# Translating Membrane Geometry into Protein Function: Multifaceted Membrane Interactions of Human Atg3 Promote LC3-Phosphatidylethanolamine Conjugation during Autophagy

**DOI:** 10.1101/2022.12.23.521840

**Authors:** Yansheng Ye, Erin R. Tyndall, Van Bui, Maria C. Bewley, Guifang Wang, Xupeng Hong, Yang Shen, John M. Flanagan, Hong-Gang Wang, Fang Tian

## Abstract

Autophagosome formation is the hallmark of macroautophagy (herein referred to as autophagy) and requires the covalent conjugation of LC3 proteins (or Atg8 in yeast) to the amino headgroup of PE (phosphatidylethanolamine) lipids. Atg3 is an enzyme that catalyzes the final step of this reaction by transferring LC3 from an LC3-Atg3 intermediate to PEs in targeted membranes. Here, we determine the solution structure of human Atg3 (hAtg3) and demonstrate that the catalytically important regions of hAtg3 are conformationally dynamic. Furthermore, we reveal that these regions and hAtg3’s N-terminal membrane curvature-sensing amphipathic helix concurrently interact with the membrane. These structural studies indicate that hAtg3 exploits a multifaceted membrane-association mechanism to position its catalytic center at the membrane surface and to bring the reaction substrates of LC3 and PE lipids to proximity for effective LC3-PE conjugation. In addition, our studies demonstrate that the interaction of the His266 residue with the membrane is primarily responsible for hAtg3’s pH-dependent activity. Our investigations advance an emerging concept that the interactions of Atg3 with the highly curved membrane rims of the phagophore spatially regulate autophagosome biogenesis.

---

Macroautophagy (autophagy) is a major intracellular degradation and recycling process that is initiated by the formation of a cup-like membrane structure called a phagophore. As the phagophore expands, cytosolic components that are targeted for degradation such as aggregated proteins, damaged organelles, and foreign organisms including viruses and bacteria, are engulfed. After the phagophore elongates, it seals and creates a double-membrane vesicle called the autophagosome. The autophagosome is then delivered to the lysosome to degrade and recycle its contents^1-5^. In eukaryotes, autophagy is essential for maintaining cellular homeostasis and basic functions; its dysfunction leads and/or contributes to many disease processes including neurodegeneration, infection, cardiovascular impairment, and tumorigenesis^6-10^.

A key step in autophagosome biogenesis is the lipidation of LC3 family proteins to phosphatidylethanolamine (PE)^5,11^. This LC3-PE conjugate regulates phagophore expansion and serves as a docking site for autophagic cargos^12-14^. This conjugate reaction is catalyzed by three ubiquitin-like conjugation enzymes: E1-like Atg7, E2-like Atg3 and the E3-like Atg12-Atg5/Atg16 complex. Despite their conceptual similarities, Atg7, Atg3, and Atg12-Atg5/Atg6 are structurally and functionally distinct from canonical E1, E2, and E3 enzymes^15-22^. For instance, *in vitro*, unlike canonical E2 enzymes, Atg3 alone selects substrate PE lipids and can catalyze the transfer of LC3 from an LC3-Atg3ß intermediate to PEs without the Atg12-Atg5/Atg16 complex^23-25^. As this reaction occurs at the membrane surface with PEs in a lipid bilayer phase, the protein-lipid interaction of Atg3 plays an essential role. In fact, Atg3’s function requires the presence of an N-terminal amphipathic helix (NAH) that selectively interacts with membranes that contain lipid-packing defects, such as the leading edge of the growing phagophore where membranes are highly curved^26-29^. Our recent studies have further revealed that the membrane association of the NAH of human Atg3 (hAtg3) is necessary, but not sufficient, to catalyze LC3-PE conjugation^30^. We have discovered a conserved region following the hAtg3 NAH that tightly coordinates its membrane geometry-sensitive interaction with its catalytic activity. In addition, enhancing the interaction of Atg3 with membranes by acetylation of its NAH reportedly extends the duration of autophagy^31^. However, how the C-terminal located catalytic center is directed to the membrane surface for effective LC3-PE conjugation remains obscure.

In this study, we have used NMR spectroscopy to determine the solution structure of hAtg3 and discovered that its NAH and catalytically important C-terminal regions concurrently interact with the membrane. The functional significance of this interaction was confirmed by *in vitro* and *in vivo* loss-of-function mutagenesis analyses. Our results suggest that hAtg3 exploits a multifaceted membrane-association mechanism^32-38^ to position the catalytic residue Cys264 at the membrane surface and bring substrates of LC3 and PE lipids in proximity to promote LC3-PE conjugation. In addition, NMR hydrogen/deuterium exchange studies revealed that the catalytic loop and following α-helix of hAtg3 are conformationally dynamic in aqueous solution and are conducive to interaction with the membrane, even though the general hAtg3 core structure is similar to those previously reported for yeast and *Arabidopsis thaliana*. Finally, we demonstrate that the interaction between hAtg3 His266 and the membrane is primarily responsible for its pH-dependent activity as reported in yeast Atg3^25^. Together, our studies strengthen an emerging concept that the interactions of Atg3 with the highly curved membrane rims of the phagophore spatially regulate autophagosome biogenesis^29,39^.

## Results

### Structure and conformational plasticity of hAtg3

hAtg3 is designated to as an intrinsically disordered protein since more than 1/3 of its 314 residues are in unstructured regions^40^. In an earlier work, we showed that residues 90 to 190 form an unstructured loop and that deletion of these amino acids (hAtg3Δ^90-190^) did not affect function *in vitro*^30^. To facilitate NMR structure determination, we employed a construct that lacks the first 25 residues of the N-terminal as well as residues 90 to 190 (hAtg3Δ^N25,^ Δ^90-190^). This construct produces high resolution spectra and is very similar to a yeast Atg3 (yAtg3) version that was recently selected for X-ray structure determination^41^. Residues 1 to 25 are unstructured in aqueous solution but they undergo a conformational change to form an amphipathic α-helix upon interaction with the membrane^42^. Their deletion (hAtg3Δ^N25,^ Δ^90-190^) causes minimal perturbation in the core structure as evidenced by an overlay of the TROSY spectra of hAtg3Δ^90-190^ and hAtg3Δ^N25,^ Δ^90-190^ shown in Extended Data Fig. 1.

An overlay of ten solution structures of hAtg3Δ^N25,^ Δ^90-190^ (determined with 2384 NOEs, 268 torsion angle restraints, and 557 RDCs measured in four alignment media) is shown in Fig. 1a; statistics from our NMR structure determination are summarized in Extended Data Table 1. Superpositions of hAtg3Δ^N25,^ Δ^90-190^ onto the structures of yeast and *Arabidopsis thaliana* Atg3 (yAtg3 and AtAtg3) demonstrate a conserved E2-like fold. The reported C^α^ RMSDs of hAtg3 to yAtg3 and AtAtg3 structures by Chimera are 1.7 Å and 1.5 Å, respectively. The largest deviation is observed in helix F (according to the secondary structure definition in the yAtg3^18^), which immediately follows the catalytic loop containing Cys264 (Fig. 1b). In addition, in the structure of hAtg3 determined here, helix F (residues 268 to 278) is notably less well defined with an RMSD of 1.0 Å while the rest of the structured elements converge to an overall RMSD of 0.4 Å. This is due to the use of a limited number of observable NOEs for the structure calculation and may reflect the flexibility of this helix.

**Fig. 1:**
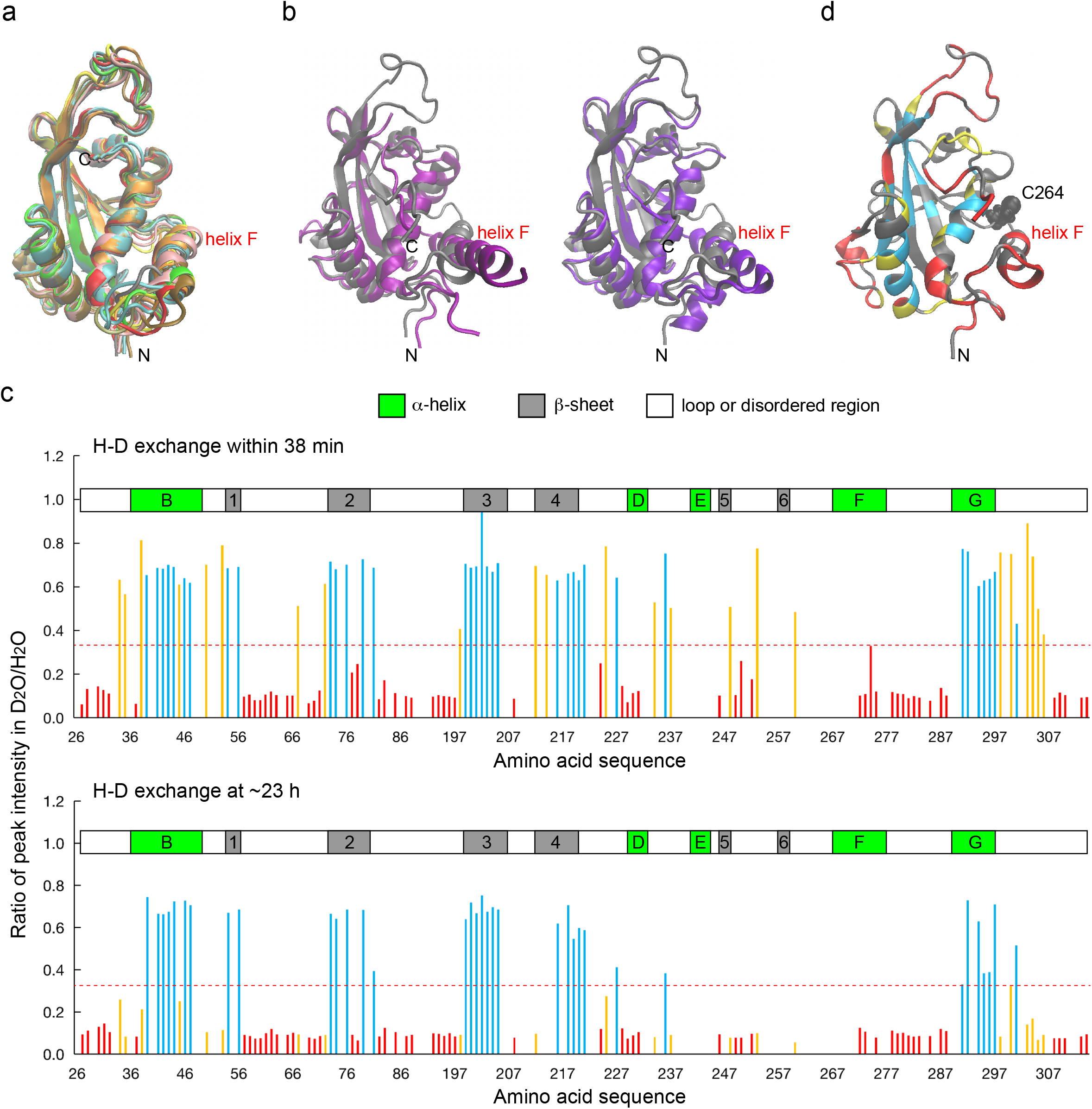
Structure and conformational flexibility of hAtg3Δ^N25,^ Δ^90-190^. **a**, Overlay of hAtg3Δ^N25,^ Δ^90-190^ ten solution structures. C-terminal unstructured residues 306 to 314 are not shown for clarity. **b**, Structural alignments of hAtg3Δ^N25,^ Δ^90-190^ (grey) with yeast Atg3 (purple, PDB 6OJJ, left), and *Arabidopsis thaliana* Atg3 (violet, PDB 3VX8, right). Helix F is indicated. **c**, H/D exchange of ^15^N-labeled hAtg3Δ^N25^, Δ^90-190^ at pH 6.5, 25 °C. Group I: residues with fast H/D exchange (more than 67% of resonance intensities lost within 38 mins after addition of D_2_O) are colored red. Group II: residues with slow H/D exchange (less than 67% of resonance intensities lost at ∼23 hrs after addition of D_2_O) colored in cyan. Group III: residues with intermediate H/D exchange (slower than Group I and faster than Group II) are colored orange. Secondary structural elements are indicated according to corresponding elements in a yeast Atg3 structure (PDB 2DYT). **d**, Residues with different H/D exchange rates are mapped onto the hAtg3 NMR structure in corresponding colors as shown in **c**. Residues with uncharacterized H/D exchange rates are shown in grey. Helix F is indicated and catalytic residue Cys264 is shown in VDW.

To characterize protein stability and the dynamic features of hAtg3Δ^N25,^ Δ^90-190^, we performed NMR hydrogen/deuterium (H/D) exchange experiments. Instead of quantitatively determining an exchange rate constant, for the purpose of this discussion, we classified the peaks observed in TROSY spectra into three groups according to their intensities in H/D exchange experiments. Group I amides include those that are in fast exchange (colored red in Fig. 1c) and is defined by amides that lost >67% of their peak intensities in the first 2D TROSY spectrum (data collection started ∼21 mins after addition of a D_2_O buffer and took ∼17 mins to finish) relative to a reference spectrum collected in an H_2_O buffer. Group II amides exchanged slowly (colored cyan in Fig. 1c); more than 33% of their initial peak intensities remained in the last 2D TROSY spectrum taken after ∼23 hrs of H/D exchange. Group III amides displayed intermediate H/D exchanges, slower than Group I and faster than Group II (colored orange in Fig. 1c). Most group I amides are located in unstructured loops but surprisingly, the amides of residues in helix F also exhibit fast exchange. Group II amides are generally located in β-sheets and helices B (residues 36 to 49) and G (residues 289 to 297). Group III amides are distributed throughout the structure. These results indicate that helix F is substantially less stable than other two long helices B and G. In addition, as we reported previously, the catalytic loop (residues 262 to 267) is likely involved in motions on the millisecond timescale since its NH peaks were not observed in the TROSY spectrum due to exchange broadening^30^. Taken together, our NMR results provide direct evidence that the catalytic loop and adjacent regions of hAtg3 are highly dynamic in aqueous solution. As we describe below, the structural changes in these regions are observed upon direct interaction with the membrane.

### Multifaceted membrane association of hAtg3

The LC3-PE conjugation reaction requires that PEs are anchored in lipid bilayers. However, the mechanism by which the catalytic site Cys264 of hAtg3 colocalizes to the bilayer surface to facilitate the transfer of LC3 to PEs remains unknown. In a recent study, we observed substantial chemical shift perturbations (CSPs) around the catalytic site of hAtg3 upon the binding of its NAH to bicelles (as we demonstrated previously, bicelles support the conjugase activity of hAtg3)^30^. Despite extensive effort, we could assign only 155 out of 199 non-proline residues of the bicelle-bound hAtg3Δ^90-190^; 22 out of 38 residues from S260 to V297 (including residues in the catalytic loop, helices F and G) remain unassigned^42^. Resonances from these residues are either missing or displayed pronounced exchange broadening in the TROSY spectrum. Thus, these regions are likely involved in conformational exchanges in bicelle-bound hAtg3Δ^90-190^. During sample optimization we noticed that hAtg3Δ^90-190^ in bicelles is stable only for one or two days at pH 6.5, but at pH 7.5 is stable for several weeks. We hypothesized that the membrane interaction of a protonated His residue in hAtg3Δ^90-190^ contributes to its instability at lower pH. This led us to substitute His266, one of two fully conserved His residues (Extended Data Fig. 2), to Leu to mimic its non-protonated state. Like the wildtype protein, hAtg3^H266L^ shows similar conjugase activity in an *in vitro* LC3B-PE conjugation assay (Extended Data Fig. 3b). Moreover, the hAtg3Δ^90-190, H266L^ construct shows dramatically improved spectral quality in bicelles. Multiple new resonances were observed and assigned to the hAtg3 catalytic region (Extended Data Fig. 3a). Further optimizations led to an hAtg3 construct containing four mutations (H240Y, V241A, P263G, and H266L, referred to as hAtg3Δ^90-190, 4M^) that produced a high quality TROSY spectrum (Extended Data Fig. 4a) and its corresponding hAtg3^4M^ construct remained functional (Extended Data Figs. 3b and 3c). To date, we have assigned all but 13 residues of the bicelle-bound hAtg3Δ^90-190, 4M^. Unassigned residues are Met1, Gln2, Asn3, Lys27, Phe28, Ala65, Asp239, Lys242, Lys243, Cys264, Arg265, His311, and Phe312. Importantly, the majority of residues adjacent to the catalytic region, including those in helices F and G, are now assigned; this allows us to analyze their perturbations upon interacting with the membrane. Extended Data Fig. 4b plots the CSPs of hAtg3Δ^90-190, 4M^ in the absence and presence of bicelles. Consistent with our previous study of wildtype protein^30^, residues showing substantial CSPs include both NAH and C-terminal residues. This suggests that the membrane-interacting surface is likely located here.

To determine the residues in hAtg3 that directly interact with the membrane, we performed an NMR cross-saturation experiment. This experiment was initially developed to map the protein-protein interfaces and has been used to identify surface residues that are directly involved in protein/lipid interactions^43-46^. On a sample of perdeuterated ^15^N, ^2^H-hAtg3Δ^90-190, 4M^ mixed with DMPC/DMPG/DHPC bicelles in an 80% D_2_O and 20% H_2_O buffer, irradiation at 1.28 ppm (hydrophobic tail residues of lipids) with a 40 ms Gaussian pulse for 1.6 s resulted in the effective saturations of all lipid resonances, except the peak from choline methyl groups, due to spin diffusion (Extended Data Fig. 5a). The selective transfer from the saturated lipid resonances to the residues in hAtg3 that are located at the protein/lipid interface results in a reduction of their resonance intensities when compared to spectra with off-resonance saturations. Figs. 2a and 2b show the results of these cross-saturation experiments for hAtg3Δ^90-190, 4M^. As expected, residues in the NAH display the largest cross-saturation effects since they form an amphipathic helix that is inserted into the membrane. Surprisingly, two stretches of C-terminal residues also display significantly reduced intensities. Region I includes residues from 262 to 277, which are in the catalytic loop and dynamic helix F (Fig. 1c). Region II includes residues from 291 to 300; most of these reside in helix G (Fig. 1c). When mapped onto the hAtg3 structure, these residues appear to cluster around the catalytic residue Cys264 (Fig. 2c). As a control, we performed the same experiment on a sample of perdeuterated ^15^N, ^2^H-hAtg3Δ^90-190, 4M^ in an 80% D_2_O and 20% H_2_O solution; all residues exhibited few cross-saturation effects as shown in Extended Data Fig. 5b. Therefore, these cross-saturation experiments provide tangible evidence that, in addition to the NAH, the C-terminal regions I and II of hAtg3 also directly interact with the membrane.

**Fig. 2:**
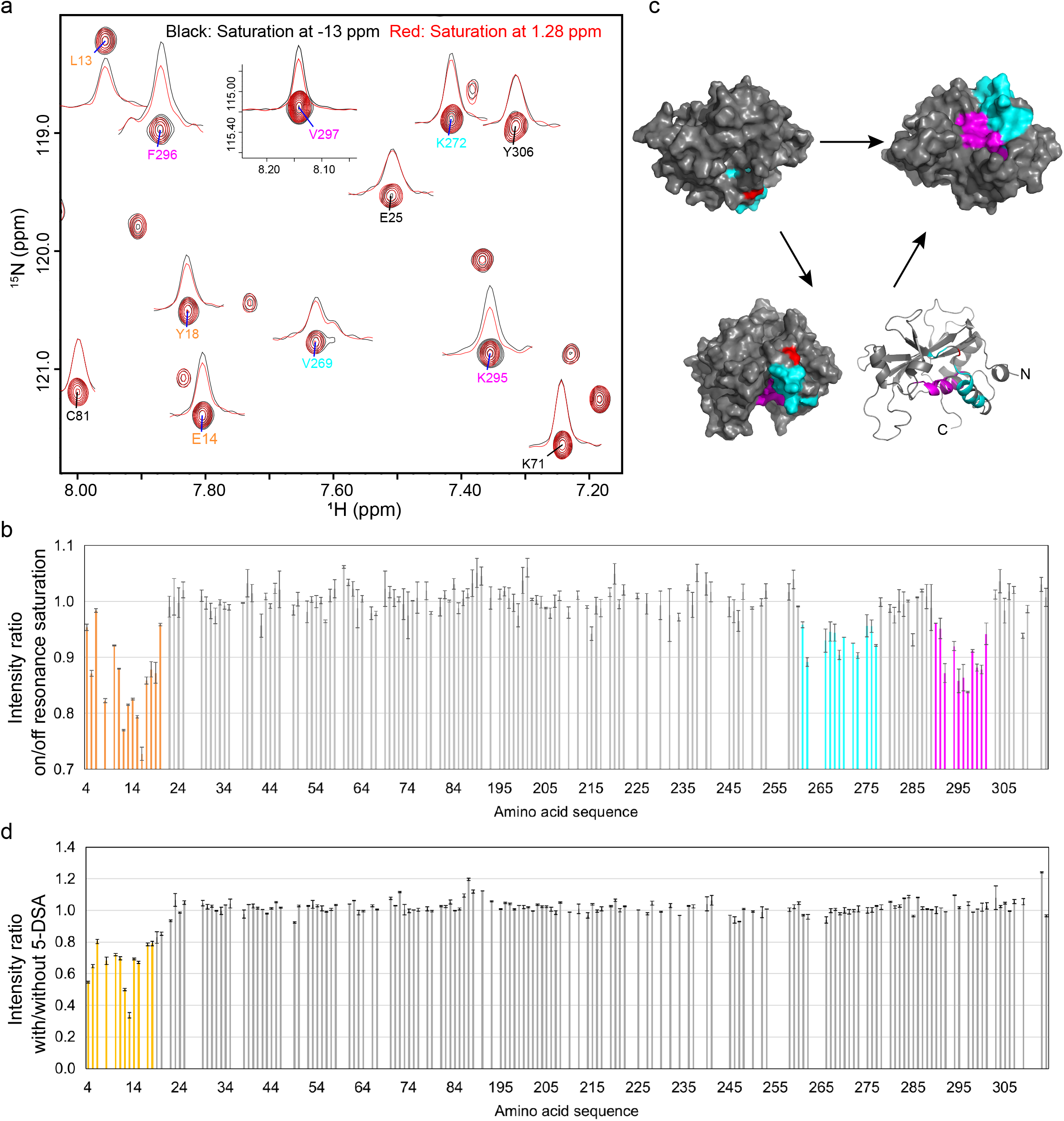
Multifaceted membrane association of hAtg3Δ^90-190, 4M^ determined by cross saturation and paramagnetic relaxation enhancement experiments. **a**, Close-up of ^1^H-^15^N TROSY spectra of perdeuterated ^15^N, ^2^H-hAtg3Δ^90-190, 4M^ in an 80% D_2_O and 20% H_2_O bicelles solution (DMPC:DMPG:DHPC = 8:2:20, q = 0.5, 12% w/v) with saturations at -13.0 ppm (black) and 1.28 ppm (red). **b**, Plot of cross saturation effects against residue numbers. The perturbed residues from the NAH are colored orange, while those from 262 to 277 and 291 to 301 are colored in cyan and magenta, respectively. **c**, Perturbed residues from 262 to 277 (cyan) and 291 to 301 (magenta) are mapped onto hAtg3Δ^N25,^ Δ^90-190^ NMR structure. Catalytic site Cys264 is indicated in red. **d**, Plot of PRE effects from 5-DSA against residue numbers. A final concentration of 1.5 mM 5-DSA was included in the NMR sample of ^15^N, ^2^H-hAtg3Δ^90-190, 4M^ in 12% (w/v) bicelles (DMPC:DMPG:DHPC=8:2:20, q=0.5).

To probe the insertion depth of these membrane-interacting regions, we performed NMR paramagnetic relaxation enhancement (PRE) experiments using 5-doxylstearic acid (5-DSA)^47-50^, a membrane soluble probe. As shown in Fig. 2d, residues in the hAtg3 NAH exhibit large PREs while all other residues experience few or no effects. Since the PRE probe is embedded in the acyl chains of the lipid bilayer, these results indicate that the NAH inserts into the hydrophobic core of the membrane. Together, our NMR chemical shift perturbation, cross-saturation, and PRE experiments demonstrate that hAtg3 interacts with the membrane using multiple surfaces. In addition to the NAH that inserts into the membrane and interacts with the hydrophobic core, we postulate that C-terminal regions I and II interact with the polar headgroup layer of the membrane.

### Mutagenesis analyses of membrane-interacting regions I and II of hAtg3

We next examined the nature of protein-lipid interactions of the two C-terminal regions identified in the cross-saturation experiment. Residues 262 to 277 of region I are highly conserved (Fig. 3a), and secondary chemical shift analyses of their assigned ^13^C^α^ and ^13^C^β^ shifts in bicelle-bound hAtg3Δ^90-190, 4M^ indicate that residues 265 to 277 are in a helical conformation (Extended Data Fig. 6). Interestingly, when plotted on a helix wheel these residues segregate into a hydrophobic or a hydrophilic face (Fig. 3a). The polar face includes residues Arg265, Glu268, Lys271, Lys272, Glu275, and Thr276 and the non-polar face includes residues His266, Ala267, Val269, Met270, Ile273, Ile274, and Val277. Thus, residues 265 to 277 form an amphipathic helix upon membrane binding. Notably, some of these hydrophobic amino acids are exposed to solvent in the apo structure of hAtg3 and are presumably primed to interact with the membrane. Consistent with this hypothesis, as shown in Fig. 3b, substitution of these hydrophobic residues with a negatively charged Asp abolishes (H266D, A267D, and I273D) or markedly reduces (V269D and M270D) LC3B-PE formation in an *in vitro* conjugation assay. The I274D mutant showed modest reduction in LC3B-PE formation, likely because Ile274 locates at the border of hydrophobic and hydrophilic faces. In addition, the substitution of positively charged Arg or Lys residues that presumably locate at the membrane/solution interface to a negatively charged Asp (R265D and K271D) also greatly reduces or disrupts the formation of LC3B-PE. As a control, we confirmed that these mutants form the LC3B-hAtg3 intermediate normally (Extended Data Fig. 7a).

**Fig. 3:**
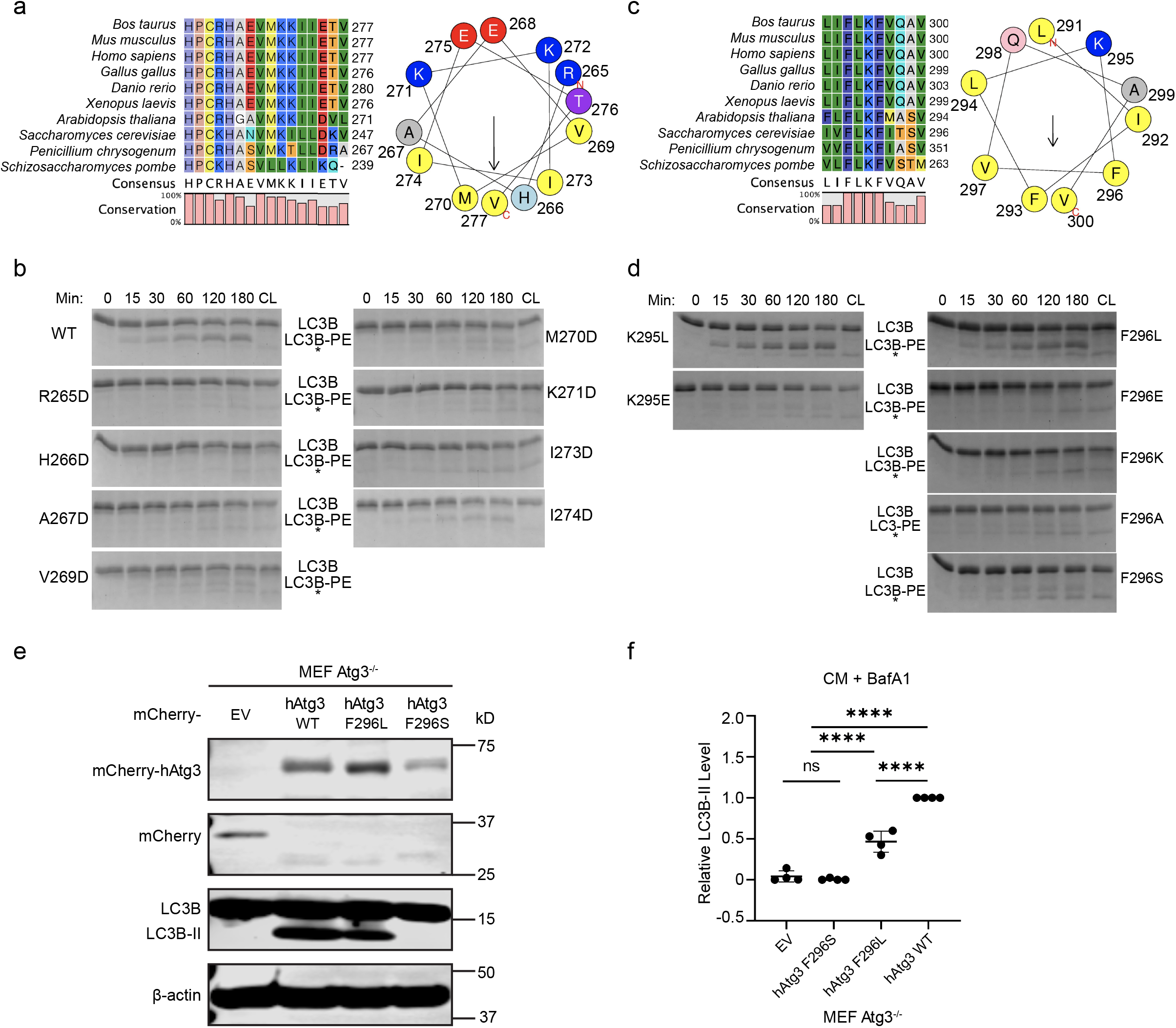
hAtg3 C-terminal interaction with the membrane is indispensable for LC3B lipidation. **a**, Left panel: sequence comparison of hAtg3 C-terminal residues 262 to 277 with the analogous region in selected organisms. Right panel: helical wheel plot for hAtg3 residues 265 to 277. **b**, Representative SDS-PAGE gel images of time dependent formation of LC3B-PE for hAtg3 and its mutants (n = 3). CL is the control without liposomes. Asterisk indicates a small amount of degradation of LC3B in the presence of ATP. **c**, Left panel: sequence comparison of hAtg3 C-terminal residues 291 to 300 with the analogous region in selected organisms. Right panel: helical wheel plot for hAtg3 residues 291 to 300. **d**, Representative SDS-PAGE gel images of time dependent formation of LC3B-PE for hAtg3 and its mutants (n = 3). CL is the control without liposomes. Asterisk indicates a small amount of degradation of LC3B in the presence of ATP. **e**, Representative immunoblot (n = 4 blots). Atg3 knockout (Atg3^-/-^) mouse embryonic fibroblasts (MEFs) stably expressing mCherry EV (empty vector), mCherry-hAtg3 WT (wildtype), mCherry-hAtg3^F296L^, or mCherry-hAtg3^F296S^ mutant were cultured in complete media (CM) with 100 nM bafilomycin A1 (BafA1 to block LC3-II degradation) for 3 hrs and subjected to immunoblotting with the indicated antibodies. **f**, Quantitative analysis of the relative LC3B–II level (n = 4 blots) in *in vivo* LC3B lipidation experiments. Statistical analysis was performed using one-way ANOVA test followed by Turkey’s multiple comparisons test. Data are presented as mean ± SD. *P* values: \*\*\*\**P* < 0.0001; ns, not significant.

For residues 291 to 300 of region II, secondary chemical shift analyses of their assigned ^13^C_α_ and ^13^C_β_ shifts in bicelle-bound hAtg3Δ^90-190, 4M^ indicate that these residues adopt a helical conformation (Extended Data Fig. 6), as in its apo structure. This helix is comprised of predominantly hydrophobic residues; a helix wheel plot shows a very large non-polar face and a small polar face containing only Lys295 and Gln298 residues (Fig. 3c). In region II, there are four consecutive residues (F^293^LKF^296^) that are fully conserved and experience large cross-saturation effects (Figs. 2a and 2b). Leu294 interacts with helix B and strands of β-sheets, and its mutations would likely disrupt hAtg3’s structure. However, we constructed single site substitutions to examine the interactions of Phe293, Lys295, and Phe296 with the membrane. The F293S mutation was not stable and was not investigated further. To test whether Lys295 is embedded into the membrane while extending or “snorkeling” its positively charged sidechain to the negatively charged phospholipid head groups, we substituted Lys295 with a Leu or a Glu residue. The hAtg3^K295L^ mutant remained fully functional, but hAtg3^K295E^ nearly completely lost its conjugase activity (Fig. 3d). Similarly, while the hAtg3^F296L^ mutant retained its enzymatic activity, replacing Phe296 with an acidic Glu (F296E), a polar Ser (F296S), or a small hydrophobic Ala (F296A) residue nearly abolished its activity. Again, these mutations form the LC3B-hAtg3 intermediate and bind the membrane normally (Extended Data Figs. 7a and 8). Thus, our mutagenesis results clearly support the association of two C-terminal regions of hAtg3 with the membrane.

The functional importance of Phe296 was recognized in a previous study of yAtg3^51^. Substitution of the corresponding Phe residue with a Ser (F293S) in yAtg3 was reported to dramatically increase its conjugase activity. This result is inconsistent with our finding that hAtg3^F296S^ is impaired in LC3B-PE conjugation, suggesting that yAtg3 and hAtg3 may have subtly different catalytic mechanisms. To investigate this further, we introduced wild-type hAtg3^WT^ and the hAtg3^F296S^ or hAtg3^F296L^ variant into Atg3^−/−^ mouse embryonic fibroblasts (MEFs) by lentiviral transduction and examined the LC3B-PE conjugation *in vivo* by western blot analysis. While the LC3B-PE conjugate (LC3B-II) was absent in control Atg3^−/−^ MEFs transduced with empty vector (EV), the expression of hAtg3^WT^ rescued the LC3B-PE conjugation (Figs. 3e and 3f). Consistent with our *in vitro* results (Fig. 3d), the introduction of the hAtg3^F296S^ variant into Atg3^−/−^ MEFs failed to rescue LC3B-II production, whereas the hAtg3^F296L^ mutant retained its ability to induce LC3B lipidation, albeit to a lesser extent than hAtg3^WT^ (Figs. 3e and 3f).

### hAtg3 His266 functions as a pH sensor

Our observed interaction between the hAtg3 C-terminal region I and the membrane requires His266 to be in a neutral nonprotonated state (Fig. 3). By extension, at low pH the protonated state of His266 should disrupt this interaction and hAtg3 should be rendered non-functional. Indeed, like yAtg3^25^, Atg3 displays maximum activity at high pH (>8.5), but its activity is markedly reduced at pH 6.5 and the protein becomes nearly inactive at pH 6.0 (Figs. 4a and 4b). When His266 is replaced by a Leu or Phe to mimic its non-protonated state, the pH-dependent activity becomes much less pronounced (Figs. 4a and 4b). Furthermore, substituting His266 with a Lys to mimic its protonated state (hAtg3^H266K^) led to a complete loss of hAtg3 conjugase activity at pH 7.5 in an *in vitro* conjugation assay (Extended Data Figs. 3b and 3c). Consistently, an overlay of TROSY spectra of hAtg3^H266K^ and hAtg3^H266L^ in bicelles indicates that they differ in how they interact with the membrane (Extended Data Fig. 3e). Therefore, we conclude that the His266 in hAtg3 is largely responsible for its pH-dependent activity and functions as a pH “switch” that regulates the interactions of regions I with the membrane.

**Fig. 4:**
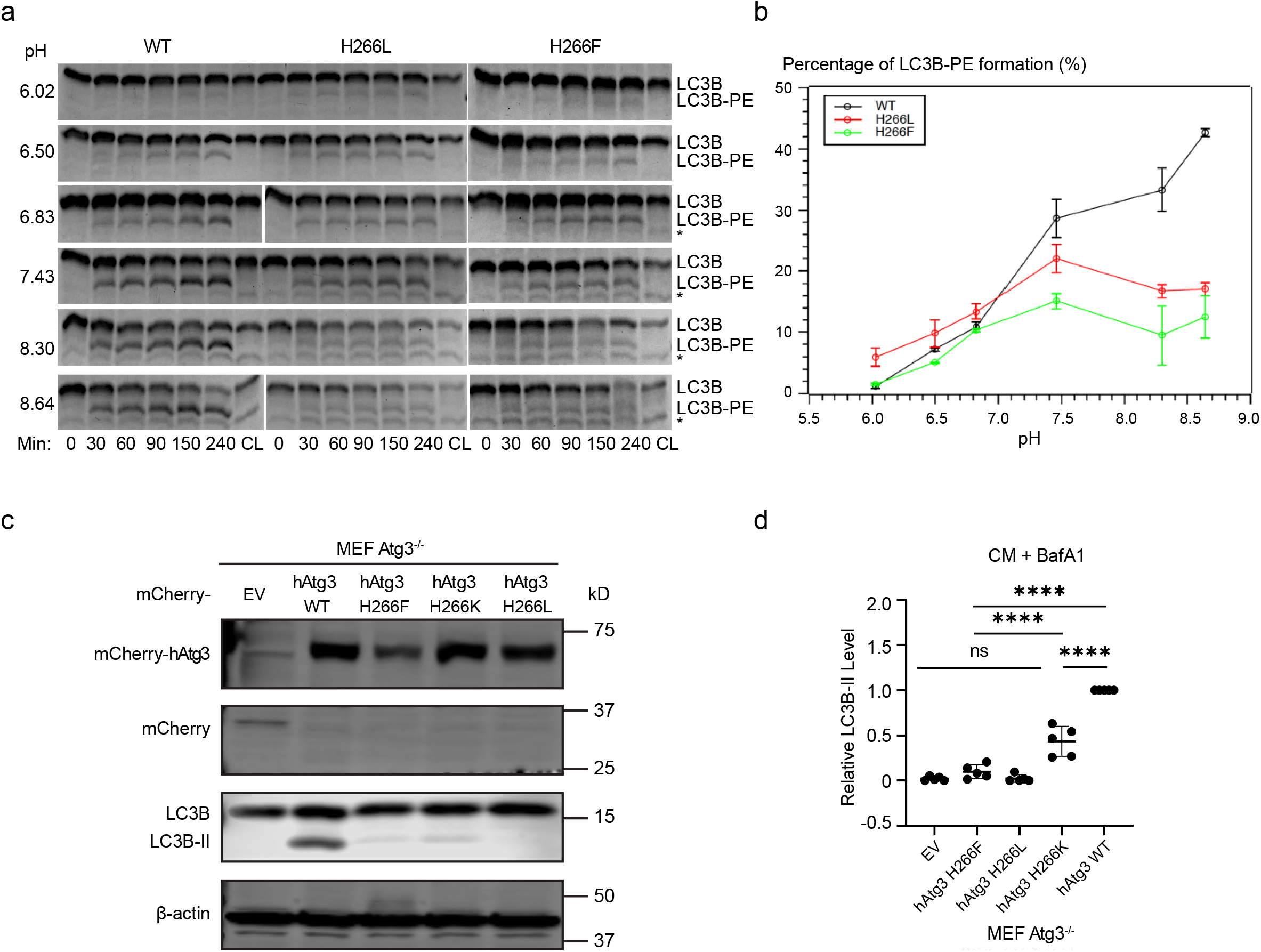
His266 is primarily responsible for hAtg3 pH-dependent activity. **a**, Representative SDS-PAGE gel images of time dependent formation of LC3B-PE for hAtg3 and its H266L and H266F mutants under different pHs (n = 3). CL is the control without liposomes. Asterisk indicates the degradation of LC3B in the presence of ATP. **b**, Plot of LC3B-PE formation at 1.5 hr for hAtg3 and its H266L and H266F mutants against pHs. Data are presented as mean ± SD. Quantification of conjugation reactions were obtained from three separate measurements (n = 3). **c**, Representative immunoblot (n = 5 blots). Atg3 knockout (Atg3^-/-^) mouse embryonic fibroblasts (MEFs) stably expressing mCherry EV (empty vector), mCherry-hAtg3 WT (wildtype), mCherry-hAtg3^H266F^, mCherry-hAtg3^H266K^, or mCherry-hAtg3^H266L^ mutant were cultured in complete media (CM) with 100 nM bafilomycin A1 (BafA1 to block LC3-II degradation) for 3 hrs and subjected to immunoblotting with the indicated antibodies. **d**, Quantitative analysis of the relative LC3B-II level (n = 5 blots) in *in vivo* LC3B lipidation experiments. Statistical analysis was performed using one-way ANOVA test followed by Turkey’s multiple comparisons test. Data are presented as mean ± SD. *P* values: \*\*\*\**P* < 0.0001; ns, not significant.

To characterize the role(s) of several other histidine residues, we performed mutagenesis studies of His240 and His287 that locate on the same side of the hAtg3 membrane-interacting interfaces identified above, and their effects were very different from the His266 mutations (Extended Data Fig. 7b). Substitutions of His240 and His287 with Lys or Leu remained functional. In addition, we substituted the fully conserved His262 with Lys, Leu, Phe, Ala, Asn, Asp, or Tyr residues. All mutants abolished LC3B-PE production (Extended Data Fig. 7c) even though they formed LC3B-Atg3 intermediates normally (Extended Data Fig. 7d). This is consistent with a previous report that the corresponding His232 residue in yAtg3 plays a critical catalytic or structural role^16^.

To characterize the role of hAtg3 His266 residue in the LC3B-PE conjugation reaction *in vivo*, we expressed the hAtg3^H266L^, hAtg3^H266K^ and hAtg3^H266F^ mutants in Atg3^−/−^ MEFs and measured the lipidated form of LC3B (LC3B-II) by western blot analysis. While the introduction of hAtg3^H266F^ and hAtg3^H266L^ into Atg3^-/-^ MEFs nearly completely failed to rescue the LC3B-PE conjugation, the expression of hAtg3^H266K^ increased LC3B-II synthesis but to a much lesser extent than hAtg3^WT^ (Figs. 4c and 4d). Collectively, these data strongly support the concept that the hAtg3 His266 residue is essential for protein function *in vivo* and cannot be replaced by an amino acid with similar hydrophobic, aromatic or hydrogen-bonding characteristics. Although we were able to mimic the interaction of His266 with the membrane using Leu and Phe in a liposome based single turnover assay *in vitro*, His266 may also serve additional functions *in vivo*, such as product release or substrate reloading that cannot be mimicked by these two amino acids.

## Discussion

During autophagy the Atg3 enzyme catalyzes a covalent conjugation of Atg8 family proteins to the amino group of PE lipids. The reaction product Atg8-PE is required for the successful formation of an autophagosome. Since this reaction is carried out at the membrane surface with PE serving as one of two reaction substrates, the functional importance of the interaction between Atg3 and membrane has been long recognized. In an early study of yAtg3, PE lipids were reportedly required to anchor in a bilayer structure for Atg8-PE conjugation to occur, and the conjugation reaction required the presence of negatively charged lipids and proceeded more efficiently with a higher percentage of PEs in the membrane (up to 75% percent)^24^. Similarly, we found that bilayer-like bicelles supported hAtg3 conjugase activity whereas DPC micelles did not^30^. Furthermore, the Atg3/membrane interaction mediated by its curvature-sensitive NAH has been well-established^27-29^. We recently demonstrated that the membrane binding of hAtg3 NAH is necessary but not sufficient for LC3-PE conjugation; we also uncovered a conserved region in the N-terminal of hAtg3 that works together with the NAH to couple its curvature-selective membrane binding to the conjugase activity^30^. In this study, using NMR CSPs, cross-saturation, and PRE experiments we reveal that, in addition to its NAH, the catalytically important C-terminal regions of hAtg3, including the loop that harbors the catalytic residue Cys264 and two C-terminal helices (F and G), directly interact with the membrane. The functional relevance of these interactions is demonstrated by *in vitro* lipidation and *in vivo* cellular assays. Intriguingly, the catalytic loop and helix F of hAtg3 are conformationally flexible in aqueous solution as evidenced by resonance exchange broadenings and fast H/D exchanges. This result further supports the model that they are conformationally plastic and are susceptible to structural rearrangements that are critical for their conjugase activity^52^. We also show that the interaction of His266 with the membrane is primarily responsible for the observed *in vitro* hAtg3 pH-dependent conjugase activity.

We recently reported substantial chemical shift perturbations in the C-terminal regions that surround the catalytic residue Cys264 of hAtg3 when its NAH binds to the membrane^30^. However, in our previous study, we were only able to obtain ∼78% of backbone resonance assignments of bicelle-bound hAtg3Δ^90-190^ because of exchange broadenings^42^. Here, we overcame this challenge by introducing mutations (e.g. H266L mutation) that stabilize hAtg3’s interaction with the membrane and achieved near-complete (∼93%) backbone resonance assignments for bicelle-bound hAtg3Δ^90-190, 4M^. Consistent with our previous study, perturbed residues induced by NAH binding to the membrane map to a continuous surface around residue Cys264 and are located on one side of the protein. Moreover, NMR cross-saturation experiments provided new evidence that, in addition to the NAH, the catalytic loop and two following helices (F and G) also directly interact with the membrane. These newly uncovered C-terminal/membrane interactions are functionally indispensable; mutations that disrupt these interactions abolish or dramatically reduce LC3-PE conjugation *in vitro* and *in vivo*. Importantly, these hAtg3 C-terminal/membrane interactions depend on the membrane binding of its NAH, but not vice versa. As shown in our previous study, the hAtg3^Δ90-190, V8D_V15K^ mutant does not bind to the membrane and few CSPs are observed for C-terminal residues (supplementary Figs. 5 and 15(a)^30^). By comparison, hAtg3 mutants that disrupt its C-terminal membrane interaction still bind to the membrane (Extended Data Fig. 8).

We propose a model that summarizes the key findings of this study and delineates the process of hAtg3/membrane interaction (Fig. 5). In the apo state, hAtg3 assumes an inactive conformation with an unstructured NAH and conformationally flexible catalytic loop and helix F. This structural plasticity of hAtg3 may be functionally advantageous for substrate loading and unloading. In the presence of highly curved membrane surfaces, the NAH rearranges into an amphipathic helix and inserts into the membrane; the catalytic loop and two adjacent C-terminal helices F and G concurrently locate to and interact with the membrane surface. We postulate that this multifaceted membrane association of hAtg3 mechanistically brings the substrates into proximity of the catalytic site, optimally orienting them via structural arrangements necessary for efficient LC3-PE conjugation, and functionally links hAtg3 conjugase activity to NAH curvature sensitivity so that limits LC3-PE conjugation to the highly curved rim of the phagophore. Thus conceptually, the membrane functions like an-E3 ligase for LC3B-PE conjugation. Recently, a coincident association of PX and C2 domains with the membrane was shown to be a major trigger for PI3KC2α activation^53^ and a multivalent association of the PHLEHA7 PH domain with phosphatidyl-inositol-phosphate (PIP) induced PIP clustering^54^. In addition, during the activation of phospholipase A2 (PLA2), the membrane may function as an allosteric ligand^55^.

**Fig. 5:**
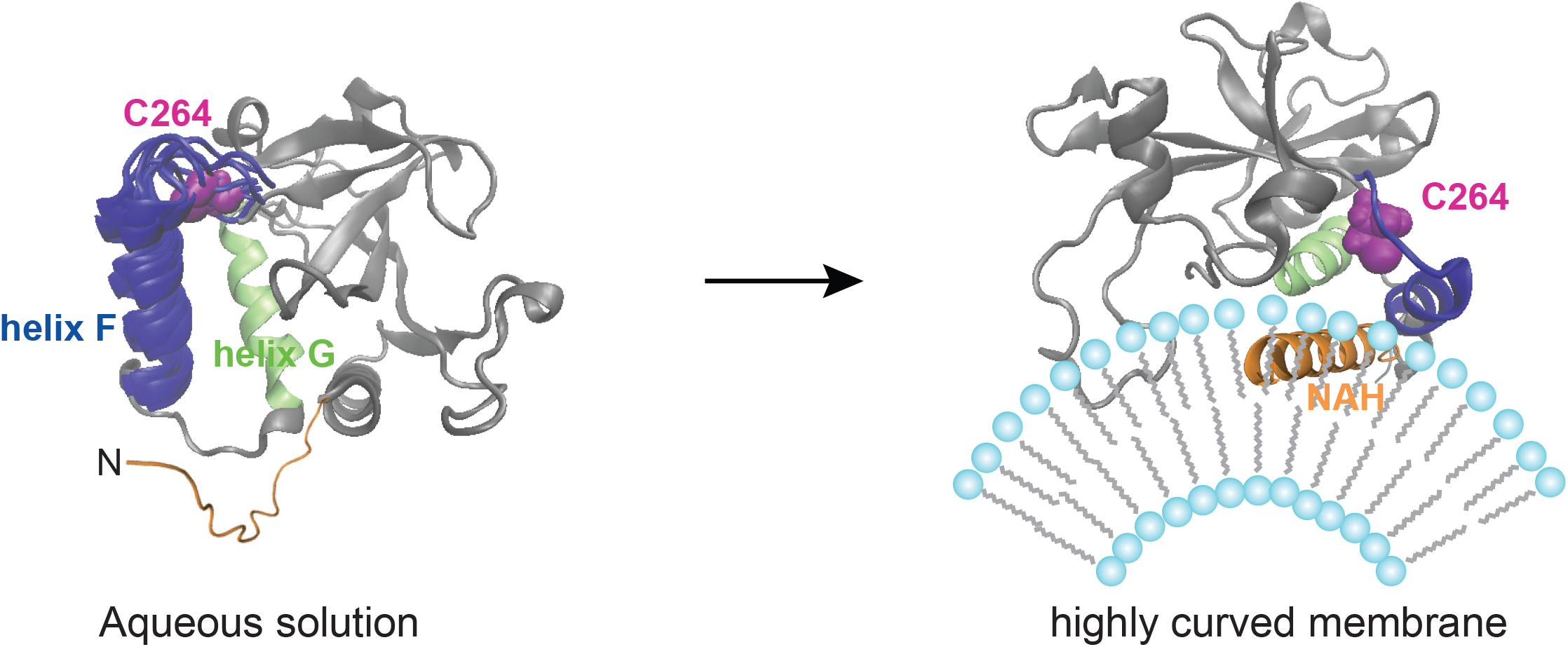
A model delineates the process of hAtg3/membrane interaction. In aqueous solution, hAtg3 has an unstructured NAH and conformationally flexible catalytic loop and helix F. At highly curved membrane surfaces, the NAH of hAtg3 rearranges into an amphipathic helix and inserts itself into the membrane; its catalytic loop and two C-terminal helices (F and G) concurrently locate to and interact with the membrane surface.

Protonation/deprotonation of a His residue is an established post-translational modification for regulating protein/protein and protein/lipid interactions^56-61^. In our model, His266 in hAtg3 locates on the membrane-interacting surface of helix F. As expected hAtg3, like yAtg3, demonstrates pH-dependent conjugase activity. Consistently, mutations that mimic the non-protonated state of His (H266L and H266F) are active and display greatly reduced pH-dependent conjugase activity, while a mutation that mimics the protonated state of His (H266K) is inactive for LC3B-PE conjugation. In addition, NMR CSPs also indicate that hAtg3 with a H266L or H266K mutation interacts differently with bicelles (Extended Data Fig. 3e). Collectively, our data support a model where His266/membrane interaction is a major regulator of the hAtg3 pH-dependent conjugase activity observed *in vitro*. Notably, the corresponding His236 residue in yAtg3 appears to function differently^51^. As reported in a previous study, the yAtg3 H236L mutant is inactive *in vitro*. By comparison, the hAtg3 H266L mutant retains ∼85% of the *in vitro* conjugase activity of the wild-type protein at pH 7.5 (Fig. 4, Extended Data Figs. 3b and 3c). This suggests that yAtg3 and hAtg3 may use different molecular mechanisms to catalyze Atg8-PE and LC3B-PE conjugations. This observation is further supported by *in vitro* and *in vivo* data which show that the hAtg3 F296S mutant abolishes LC3B-PE conjugation while the corresponding yAtg3 F293S mutant markedly increases Atg8-PE conjugation^51^.

In summary, in this study we determined the solution structure of hAtg3 and demonstrated that its catalytic loop and adjacent helix F are conformationally dynamic in aqueous solution. Furthermore, we revealed that hAtg3 associates with the membrane using multiple facets including the NAH and its catalytically important C-terminal regions. The functional implications of these hAtg3 C-terminal/membrane interactions were demonstrated by mutagenesis and *in vitro* and *in vivo* assays. Future studies, in particular the determination of membrane-bound hAtg3 and hAtg3-LC3 structures, will illustrate how these interactions brings its two substrates (PE and LC3) close into proximity and rearranges its active site to promote LC3-PE conjugation.

## Methods

### Mammalian cell culture

HEK293T cells (ATCC; CRL-3216) and Atg3^-/-^ mouse embryonic fibroblasts (MEFs) provided by Dr. Shengkan (Victor) Jin (Rutgers University -Robert Wood Johnson Medical School, NJ) were cultured in Dulbecco’s Modification of Eagle’s Medium (DMEM) supplemented with 10% fetal bovine serum and 1x Antibiotic Antimycotic Solution (AA) (Cytiva, SV30079.01).

### Plasmid construction for *in vivo* functional assay

hAtg3^WT^, hAtg3^H266L^, hAtg3^H266F^, hAtg3^H266K^, hAtg3^F296S^, and hAtg3^F296L^ cDNAs were amplified by PCR (forward: 5’-GAGCTGTACAAGTCTAGAGTGATGCAGAATGTGATTAATA-3’; reverse 5’-CGCAGATCCTTGCGGCCGCGTTACATTGTGAAGTGTCTTG-3’). Purified PCR products were sub-cloned into pCDH1-CMV-mCherry-MCS-EF1-puro viral backbone using NheI and BamHI.

### Lentiviral packaging, transduction, and cell sorting

Lentivirus was made by transfecting HEK293 cells with ViraPower packaging plasmids (pLP1, pLP2, and pLP/VSVG) from Invitrogen along with the viral vector using JetPrime. Four hours after transfection, the culture medium was replaced. Collecting the viral supernatant was done at 24 and 48 hrs post-transfection, and filtered with 0.45 μM supor membrane. The viral supernatant was added to Atg3^-/-^ MEFs and incubated for 24 hrs. Puromycin selection was started after 48 hrs post-transduction and remained for 3 days. MEF cells were sorted with flow cytometry based on mCherry expression level. MEF cells with high expression of mCherry were used for *in vivo* experiments.

### *In vivo* LC3-II synthesis measurement

Cells were incubated in complete media in the presence of 100 nM bafilomycin A1 (to block the lysosomal turnover of LC3-II) for 3 hrs and subjected to immunoblotting using the mCherry antibody (Abcam, ab125096) with a ratio of 1:1000, the β-actin antibody (Sigma, A5441-100UL) with a ratio of 1:10,000, and the LC3 antibody (Novus Biologicals, NB100-2220) with a ratio of 1:5,000. To quantify the LC3-II level, the experiments were repeated 4-5 times. The signal intensity of LC3-II was measured using Image Studio version 5 software (LI-COR Biotechnology), then normalized to the intensity of β-actin of the same sample. To calculate the relative LC3-II level, the LC3-II value of Atg3^−/−^ MEFs expressing mCherry-hAtg3^WT^ was set to be 1.

### Protein expression and purification

Full-length human Atg3 (hAtg3) was subcloned into the pET28a expression vector at the BamHI/XhoI site with His-tag and T7-tag and a thrombin cleavage site at the N-termini. All hAtg3 mutants were generated using the Q5 Site-Directed Mutagenesis Kit (New England Biolabs) with corresponding primers listed in Extended Data Table 2 and verified by sequencing. Plasmids containing target genes were transformed into chemically competent Rosetta™(DE3) pLysS cells for expression. Proteins were expressed and purified similar to previous report^30^. Typically, a single colony was selected to grow in small volume of LB medium overnight at 37 °C as a starter culture and then inoculated into a large volume of LB medium (for unlabeled proteins) or M9 medium supplemented with D-glucose (3 g/L, or D-glucose-^13^C_6_ and D_2_O) and ^15^NH_4_Cl (1 g/L) for ^15^N (or ^15^N/^13^C/^2^H, or ^15^N/^2^H) labeled samples at 37 °C until OD600 reached 0.6. The temperature was then lowered to 25 °C and cells were induced with 0.5 mM IPTG for ∼16 h and then harvested by centrifugation.

Cell pellets containing expressed proteins were suspended in a lysis buffer of 20 mM phosphate, pH 7.5, 300 mM NaCl, 2 mM beta mercaptoethanol (BME), and 1 mM MgCl_2_ supplemented with benzonase nuclease and complete protease inhibitor cocktail (Roche), and were lysed using sonication on ice with 2s on and 7s off intervals for 18 min total duration. Cell debris was removed by centrifugation (Sorvall RC5B Plus Refrigerated Centrifuge) at 26,900g at 10 °C for 30 min. Supernatants were collected, and loaded onto a Ni-NTA column (HisTrap HP). The column was washed with PBS buffers (20 mM phosphate, pH 7.5, 300 mM NaCl, 2 mM BME) without and with 30 mM imidazole, and then eluted with PBS buffer containing 500 mM imidazole. For biochemical assays, the elute of hAtg3 and its mutants (with purities no less than 90% after analysis on an SDS-PAGE gel) were concentrated. For NMR experiments, the elute of proteins from Ni-column were concentrated and exchanged to a buffer containing 50 mM HEPES, pH 7.5, 150 mM NaCl, and 2 mM BME, and followed by the addition of 0.1% (v/v) TWEEN20 and 50 units thrombin to remove T7-tag and His-tag for an overnight agitation at 4 °C. The solution was then subjected to a Q-column (HiTrap Capto™ Q) and further purified by size-exclusion chromatography using an S200 column (HiLoad 16/60 Superdex 200) and a buffer of 50 mM HEPES, 1 M NaCl, and 1 mM DTT. LC3B and mouse Atg7 for *in vitro* assays were prepared as described previously^30^. Purified proteins were exchanged into a buffer of 50 mM HEPES, pH 7.5 (for functional assays, NMR in bicelles and aqueous solution) or pH 6.5 (for NMR structure determination in aqueous solution), 150 mM NaCl, and 2 mM TCEP (tris(2-carboxyethyl)phosphine). Protein concentration was assayed using a Nanodrop (Thermo Scientific (Waltham, MA)).

### *In vitro* conjugation assay

*In vitro* LC3B-PE conjugation was conducted as previously described^30^. Typically, 5.0 μM hAtg3 or its mutant was mixed with 5 μM LC3B, 0.5 μM mouse Atg7, 1 mM MgCl_2_ in a reaction buffer (50 mM HEPES, 150 mM NaCl, 2 mM TCEP, and pH 7.5) of 37 μL. 3.7 μL were removed for a control. Then 1 mM ATP (0.45 μL of 100 mM ATP) was added and the reaction was allowed to proceed for 30 minutes at 37 °C for intermediate formation. 3.75 μL were removed to check for intermediate formations, and an additional 3.75 μL were incubated separately as a non-liposome containing control (CL). 1 mM liposomes (8.75 μL of 4 mM stock) were then added to the reaction system (final volume of 35 μL). The reaction was allowed to proceed at 37 °C, with 4.5 μL samples removed at 0, 15, 30, 60, (90), 120 (150), and 180 (240) minutes. Samples were mixed with 4 μL 4X protein loading buffer (10% w/v SDS, 10% BME, 40% Glycerol, 250 mM Tris-HCL, pH 6.8, and 0.4% bromophenol blue dye), and stored at -20 °C until analyzed by 18% polyacrylamide gel electrophoresis after heating for 3 minutes. For the gel analysis of intermediates, 4X loading buffer without BME was used. Gels were imaged on a BioRad Chemidock MP imager, and analyzed using ImageJ.

### Alignment media preparation

For the RDC experiment, we prepared charged polyacrylamide gels as described previously^62^. Briefly, stock solution of 40% neutral acrylamide and N,N’-methylenebisacrylamide in a 19:1 ratio (Bio-Rad) were mixed with a stock solution of 40% positively charged (3-acrylamidopropyl)-trimethylammonium chloride (75% wt, Aldrich) or negatively charged 2-acrylamido-2-methyl-1-propanesulfonic acid (AMPS, Sigma-Aldrich, Inc) containing N,N’-methylenebisacrylamide (Sigma) in a 19:1 ratio in equal volume. The mixtures or the stock solution containing only neutral acrylamide were diluted with 10xTBE buffer (Invitrogen) to a final 5% acrylamine concentration for the negatively-charged or neutral gel, or a final 7% acrylamine concentration for the positively-charged gel. Polymerization was initiated by the addition of 0.1% ammonium peroxide sulfate and 1% tetramethylethylenediamine (TEMED). The polymerization of 130 µL of the mixtures was carried out in a 3.2 mm diameter plastic tube. Polymerized gels were extensively washed in deionized water, and water was changed at least 3 times within 2 days. The gels were dried over a 2-day period at 37 °C on a plastic support wrapped with polyvinylidene chloride (PVDC) foil. For the NMR sample, the dried gel was transferred to a 5 mm Shigemi tube and the protein solution was added. Vertical compression of the gel was achieved by inserting the Shigemi plunger into the tube and limiting the final length at around 14 millimeters. After equilibration for at least 8 hr, the sample was ready for NMR measurements. For a phage aligned NMR sample, 500 μM ^15^N, ^2^H-labeled protein was dissolved in 50 mM HEPES, pH 6.5, 150 mM NaCl and 2 mM TCEP containing 6.5 mg/mL phage Pf1 (ASLA BIOTECH).

### NMR spectroscopy

Typical NMR samples contained 0.5 mM labeled proteins in a buffer of 50 mM HEPES, pH 6.5, 150 mM NaCl, 2 mM TCEP, and 0.02% NaN_3_. For H/D exchange experiments, 0.5 mM ^15^N-labeled proteins in 50 mM HEPES, pH 6.5, 150 mM NaCl, and 2 mM TCEP were lyophilized, and the powder was then dissolved in D_2_O (or 4% D_2_O/96% H_2_O, for reference) for immediate NMR collection. For bicelle samples, 0.5 mM ^15^N,^13^C,^2^H-labeled proteins were mixed with a final concentration of 12% (w/v) bicelles (DMPC:DMPG:DHPC = 4:1:20 (molar ratio, q=0.25), or DMPC:DMPG:DHPC = 8:2:20 (molar ratio, q=0.5, for cross saturation and PRE 5-DSA experiments) in 25 mM HEPES, pH 7.5, 150 mM NaCl, 2 mM TCEP.

All NMR data were acquired at 25 °C on a Bruker 600 MHz spectrometer except the NOESY data that were acquired on a Bruker 850 MHz spectrometer. Both instruments are equipped with cryoprobes. The data were processed using NMRPipe and analyzed using NMRView. Backbone resonance assignments were carried out using triple resonance HNCO, HN(CA)CO, HNCA, HN(CO)CA, HNCACB, and HN(CO)CACB experiments. Aliphatic and aromatic side-chain resonance assignments were obtained from the 3D ^13^C-edited HCCHTOCSY, ^13^C-edited NOESY, HCACO, ^15^N-edited NOESY and HBHA(CO)NH, 2D Hb(CbCgCd)Hd and Hb(CbCgCdCe)He spectra. The 3D ^13^C-NOESY and ^15^N-NOESY experiments were collected with 150 ms mixing time. ^1^D_NH_ and ^2^D_C’H_ RDCs for resonances with little perturbation between an aqueous solution and alignment medium were measured using the ARTSY method^63^ and an IP-HSQC experiment^64^, respectively.

Torsion angle constraints were derived from C_α_, C_β_, N, and C’ chemical shifts using TALOS+. NOESY assignments were obtained by a combined manual and automated analysis with CYANA 3.0. The structures were calculated and refined with XPLOR-NIH 3.3. PROCHECK, PSVS, and PDB validation server were used to analyze the ten lowest-energy structures with acceptable covalent-geometry.

## Data and code availability

NMR resonance assignments have been deposited with the BMRB with accession number 51749 and 51749. NMR structure has been deposited with the Protein Data Bank with accession ID 8FKM. All other data that support the findings of this study are available from the corresponding authors upon reasonable request.

## Acknowledgements

We are thankful for financial support from the National Institutes of Health NIGMS (R01 GM127730, R01 GM127954, R01 CA222349) and Four Diamonds (grants to FT and JMF). The NMR instruments used in this project were funded, in part, by the Pennsylvania State University College of Medicine via the Office of the Vice Dean of Research and Graduate Students and the Pennsylvania Department of Health using Tobacco Settlement Funds (CURE). The content is solely the responsibility of the authors and does not necessarily represent the official views of the University or College of Medicine. The Pennsylvania Department of Health specifically disclaims responsibility for any analyses, interpretations, or conclusions.

## Author contributions

Y.S.Y., E.R.T., V.B., M.C.B., G.F.W., X.H., Y.S., and F.T. designed and performed the experiments and analyzed the data. Y.S.Y., V.B., M.C.B., and F.T. drafted figures and the paper. M.C.B. and J.M.F. helped with protein preparation, data analysis, and manuscript preparation. F.T. and H.G.W. conceived, planned, and supervised this study, and wrote the final paper with feedback from all authors.

