## Supplemental Data for "Translating Membrane Geometry into Protein Function: Multifaceted Membrane Interactions of Human Atg3 Promote LC3-Phosphatidylethanolamine Conjugation during Autophagy"

**Extended Data Table 1: NMR constraints and structure statistics for hAtg3$^{\Delta N25,\ \Delta 90\text{-}190}$**
(NOE, Nuclear Overhauser Effect; RMSD, root-mean-square deviation)

| | *XPLOR-NIH* |
|---|---|
| ***NMR constraints*** | |
| *NOE distances* | 2384 |
| intra-residue | 639 |
| sequential (\|*i*-*j*\| = 1) | 642 |
| medium range (2 ≤ \|*i*-*j*\| ≤ 4) | 376 |
| long range (\|*i*-*j*\| ≥ 5) | 727 |
| *Dihedral angles* | |
| phi | 135 |
| psi | 133 |
| *Residual dipolar couplings ($^{1}D_{NH}$ and $^{2}D_{C'H}$)* | 557 |
| ***Structure statistics*** | |
| *Ensemble RMSD** | |
| backbone heavy atoms (Å) | 0.4 |
| all heavy atoms (Å) | 0.9 |
| *Ramachandran analysis*** | |
| most favored | 94.7% |
| additionally allowed | 5.3% |
| generally allowed | 0.0% |
| disallowed | 0.0% |
| *Violations (mean ± standard deviation)* | |
| distance constraints (Å) | 0.073 ± 0.003 |
| dihedral angle constraints (°) | 0.944 ± 0.172 |
| Residual dipolar couplings (Hz) | 0.716 ± 0.025 |
| *Deviations from idealized geometry* | |
| bond lengths (Å) | 0.004 ± 0.000 |
| bond angles (°) | 0.707 ± 0.027 |
| improper (°) | 0.614 ± 0.019 |

* Evaluated for secondary structure elements (excluding residues 268-278): 36-49, 54-56, 73-80, 199-207, 212-219, 229-232, 241-244, 245-248, 258-261, 289-297.

** Evaluated by Procheck for secondary structure elements (excluding residues 268-278): 36-49, 54-56, 73-80, 199-207, 212-219, 229-232, 241-244, 245-248, 258-261, 289-297.

**Extended Data Table 2: Primers (F, Forward; R, Reverse) used in this study.**

| Primers ID for in vitro experiments | | Sequences from 5' to 3' |
|---|---|---|
| H266L | F | CCATGCAGGCTTGCTGAGGTG |
| | R | GTGAACTGAACACATGGGAG |
| H266K | F | CCCATGCAGGAAAGCTGAGGTGA |
| | R | TGAACTGAACACATGGGAGG |
| H266F | F | TGAACTGAACACATGGGAGG |
| | R | TGAACTGAACACATGGGAG |
| H240Y/V241A | F | CAGTCAGGATTATGCGAAGAAAACAG |
| | R | ATGTCTTCATACATGTGC |
| P263G/H266L | F | AGGCTTGCTGAGGTGATGAAGAAAATC |
| | R | GCATCCGTGAACTGAACACATGGG |
| H240K | F | CAGTCAGGATAAAGTGAAGAAAACAG |
| | R | ATGTCTTCATACATGTGC |
| H240L | F | AGTCAGGATCTTGTGAAGAAAAC |
| | R | GATGTCTTCATACATGTGC |
| H287L | F | ACTTGGAGTTCTCATGTATCTTCTTATTTTCTTGAAATTTG |
| H287L/K | R | TCTCCCCCTCCTTCTGCA |
| H287K | F | ACTTGGAGTTAAAATGTATCTTCTTATTTTCTTGAAATTTG |
| K295E | F | TATTTTCTTGGAATTTGTACAAGC |
| K295E/L | R | AGAAGATACATATGAACTCC |
| K295L | F | TATTTTCTTGCTATTTGTACAAGCTGTC |
| F296L | F | TCTTGAAATTAGTACAAGCTGTC |
| | R | AAATAAGAAGATACATATGAACTC |
| F296K | F | TTTCTTGAAAAAAGTACAAGCTGTCATTC |
| F296E | F | TTTCTTGAAAGAAGTACAAGCTGTCATTC |
| F296A | F | TTTCTTGAAAGCTGTACAAGCTGTCATTC |
| F296K/E/A | R | ATAAGAAGATACATATGAACTCC |
| F296S | F | TTCTTGAAATCTGTACAAGCTG |
| | R | AATAAGAAGATACATATGAACTC |
| I274D | F | GAAGAAAATCGATGAGACTGTTGCAGAAGG |
| | R | ATCACCTCAGCATGCCTG |

| | | |
|---|---|---|
| I273D | F | GATGAAGAAAGACATTGAGACTGTTGCAGAAGG |
| | R | ACCTCAGCATGCCTGCAT |
| K271D | F | TGAGGTGATGGACAAAATCATTGAGACTGTTGC |
| | R | GCATGCCTGCATGGGTGA |
| M270D | F | TGCTGAGGTGGATAAGAAAATCATTGAGACTGTTGCAGAAG |
| | R | TGCCTGCATGGGTGAACT |
| V269D | F | CATGCTGAGGACATGAAGAAAATCATTG |
| | R | CCTGCATGGGTGAACTGA |
| A267D | F | TGCCTGCATGGGTGAACT |
| | R | TGGGTGAACTGAACACATG |
| H266D | F | CCCATGCAGGGATGCTGAGGT |
| | R | TGAACTGAACACATGGGAGGTG |
| R265D | F | TGAACTGAACACATGGGAGGTG |
| | R | ACTGAACACATGGGAGGTG |
| H262A | F | GTGTTCAGTTGCCCCATGCAGGCATG |
| | R | ATGGGAGGTGGTGGCAGA |
| H262N | F | GTGTTCAGTTAACCCATGCAGG |
| | R | ATGGGAGGTGGTGGCAGA |
| H262K | F | GTGTTCAGTTAAGCCATGCAGGC |
| | R | ATGGGAGGTGGTGGCAGA |
| H262D | F | GTGTTCAGTTGACCCATGCAGG |
| | R | ATGGGAGGTGGTGGCAGA |
| H262L | F | TGTTCAGTTCTCCCATGCAGGC |
| | R | CATGGGAGGTGGTGGCAG |
| H262Y | F | GTGTTCAGTTTACCCATGCAGGC |
| | R | ATGGGAGGTGGTGGCAGA |
| H262F | F | GTGTTCAGTTTTCCCATGCAGGCATG |
| | R | ATGGGAGGTGGTGGCAGA |

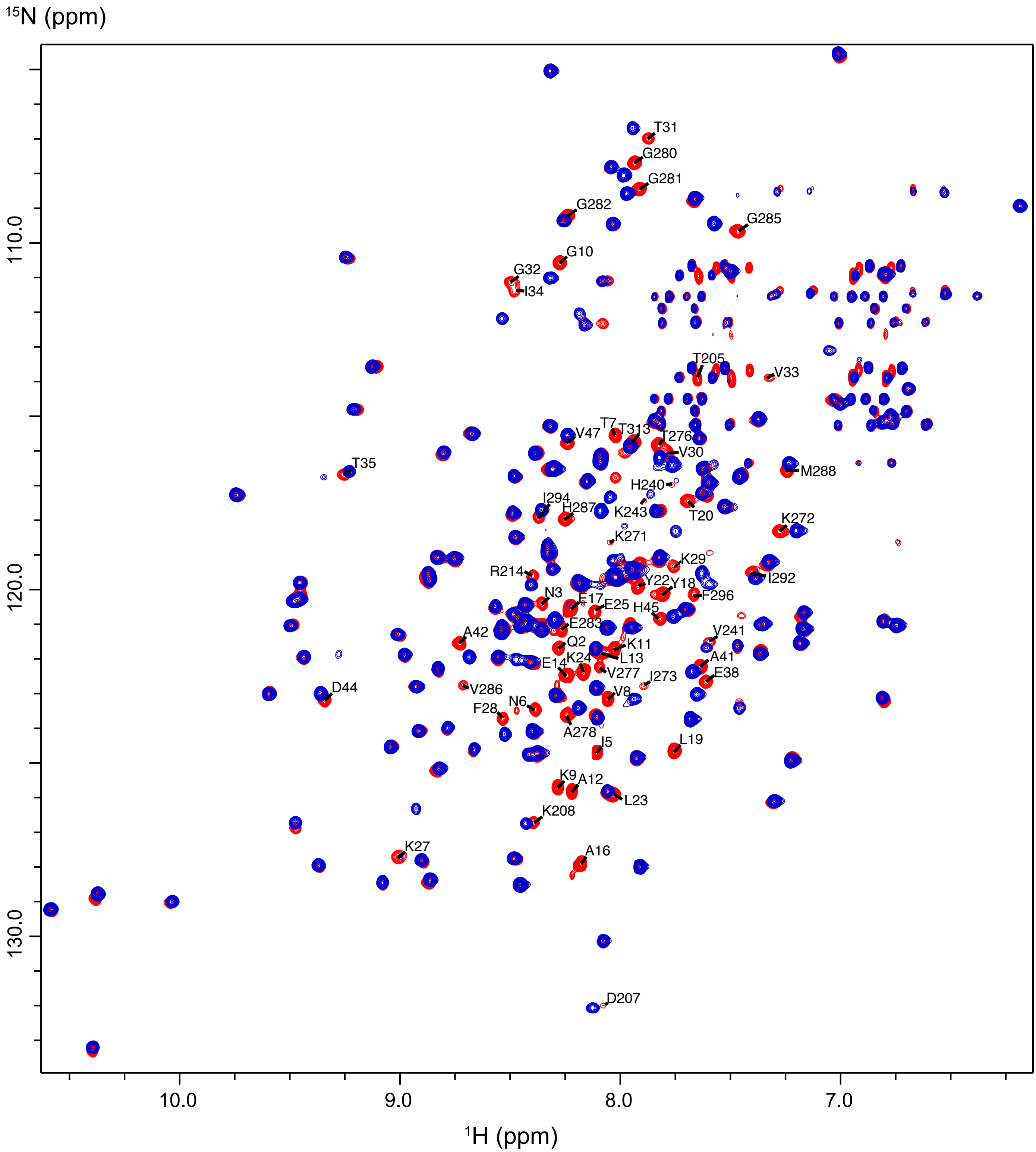


**Extended Data Fig. 1:** Deletion of the first 25 residues of the N-terminal of hAtg3 causes minimal perturbations to hAtg3's core structure, although some perturbations of C-terminal residues are observed.

Overlay of $^{15}N$-$^{1}H$ TROSY spectra of $^{15}N$-labeled hAtg3$^{\Delta 90-190}$ (red) and hAtg3$^{\Delta N25,\ \Delta 90-190}$ (blue) at pH 6.5. Perturbed residues are indicated.

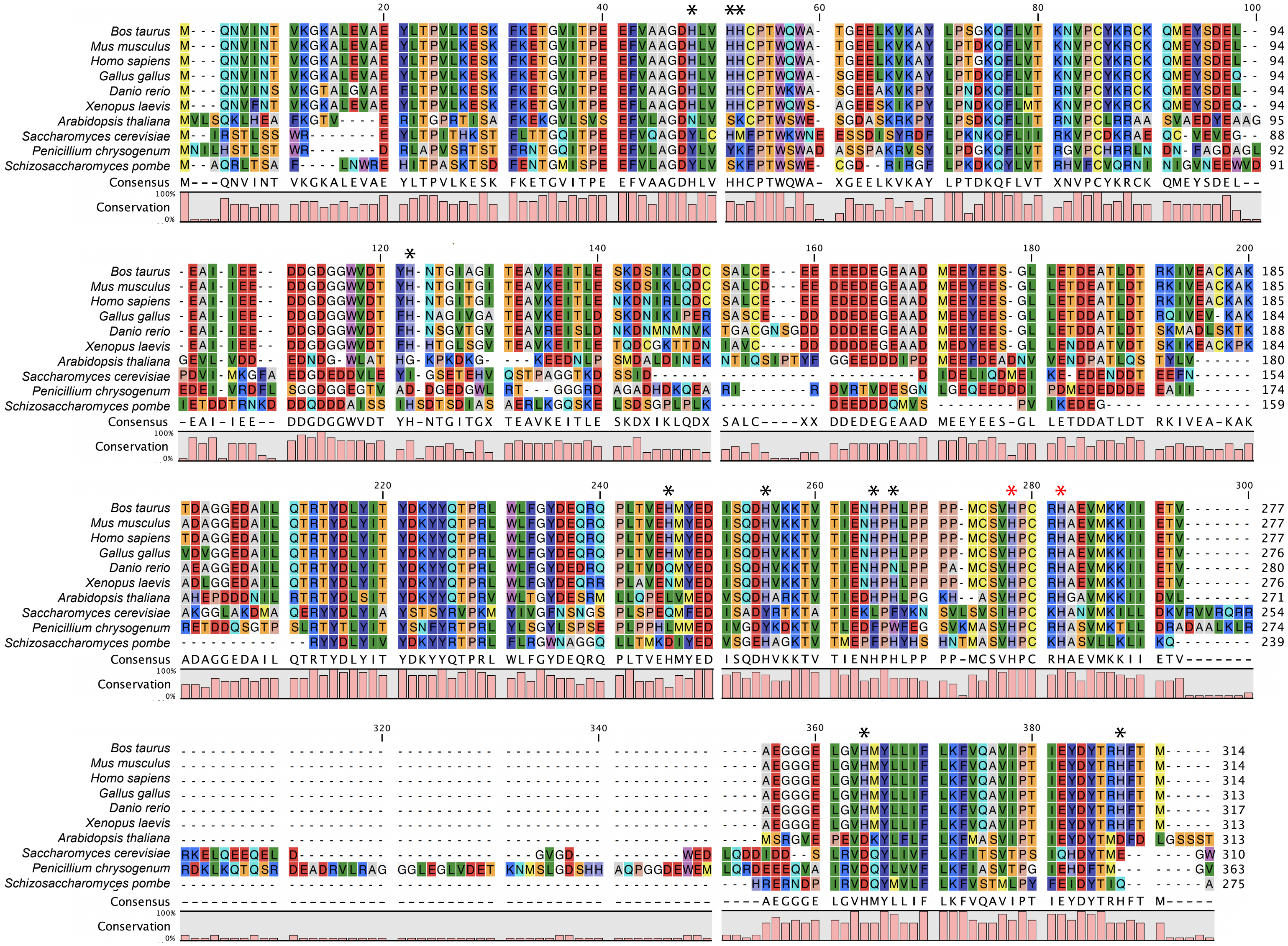


**Extended Data Fig. 2:** Sequence alignment of Atg3 homologs.
His residues of human Atg3 are indicated by asterisks (red asterisks for conserved His).

**Extended Fig. 3:** H266L mutation stabilizes hAtg3 interaction with bilayer-like bicelles.

**(a)** Overlay of $^{2}$H,$^{15}$N,$^{13}$C-labeled hAtg3$^{\Delta 90\text{-}190,\ H266L}$ (black) and hAtg3$^{\Delta 90\text{-}190}$ (red) TROSY spectra in bicelles (DMPC:DMPG:DHPC = 4:1:20, molar ratio). Multiple new resonances in the spectrum of hAtg3$^{\Delta 90\text{-}190,\ H266L}$ were observed and assigned to the hAtg3 catalytic region. Several perturbed residues are circled.

**(b)** SDS-PAGE gel images of time-dependent formation of LC3B–PE for hAtg3 (WT) and H266L, H266K, H240Y/V241A/P263G/H266L (referred to as 4M) mutants. CL represents the control without liposomes. Asterisk indicates a small amount of degradation of LC3B in the presence of ATP.

**(c)** Plots of time-dependent formation of LC3B–PE for hAtg3 and mutants. Data are presented as mean ± SD. Quantification of conjugation reactions was obtained from three separate measurements (n= 3).

**(d)** Overlay of $^{2}$H,$^{15}$N,$^{13}$C-labeled hAtg3$^{\Delta 90\text{-}190,\ H266K}$ (black) and hAtg3$^{\Delta 90\text{-}190}$ (red) TROSY spectra at pH 7.5 in bicelles (DMPC:DMPG:DHPC = 4:1:20, molar ratio). Multiple new resonances in the spectrum of hAtg3$^{\Delta 90\text{-}190,\ H266K}$ were observed and labeled with their assignments. Several perturbed resonances are circled; perturbed resonances from N-terminal residues are indicated in green.

**(e)** Overlay of $^{2}$H,$^{15}$N,$^{13}$C-labeled hAtg3$^{\Delta 90\text{-}190,\ H266K}$ (black) and hAtg3$^{\Delta 90\text{-}190,\ H266L}$ (red) TROSY spectra at pH 7.5 in bicelles (DMPC:DMPG:DHPC = 4:1:20, molar ratio). Perturbed resonances are indicated; N-terminal residues are shown in green.

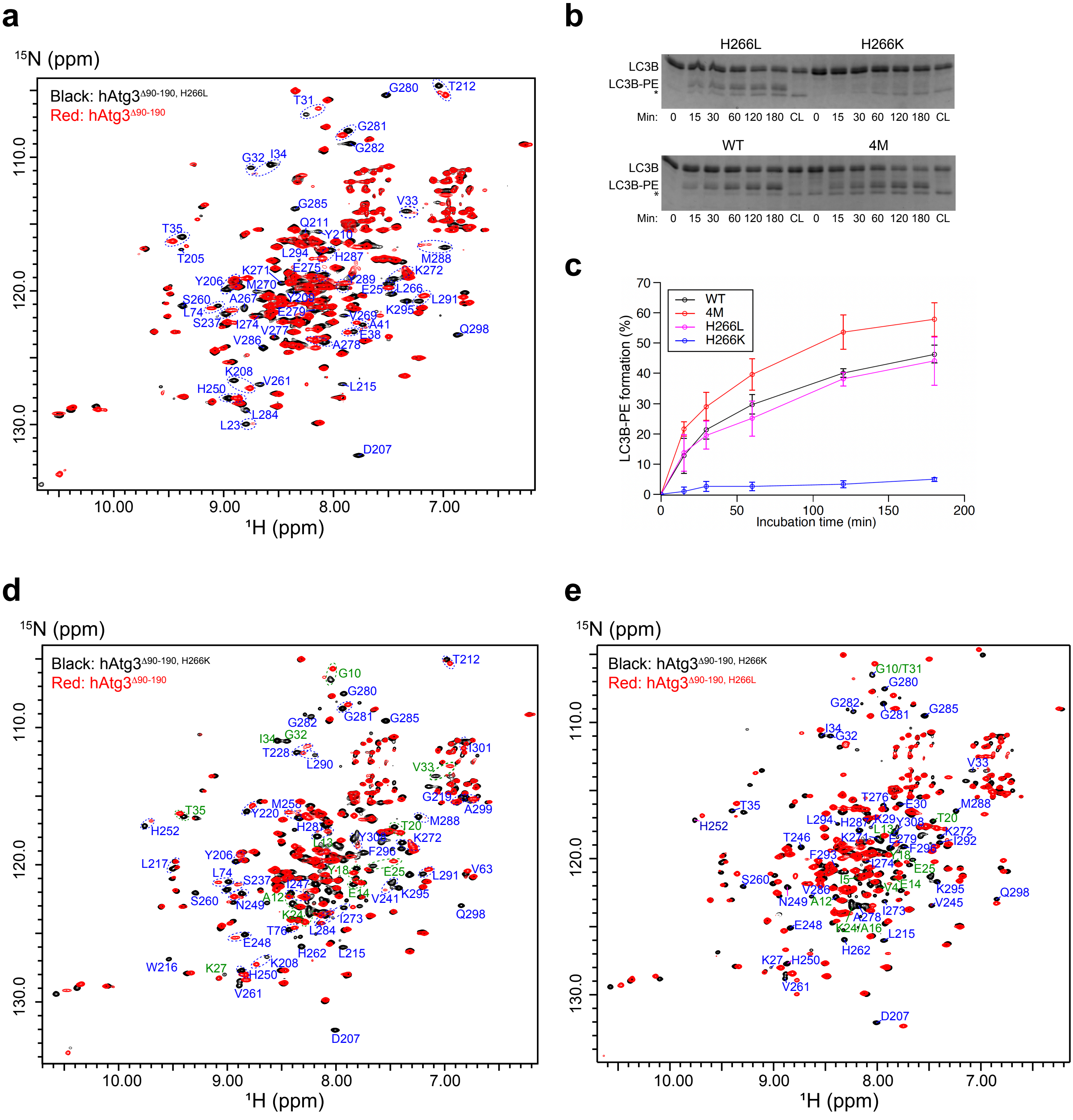


**Extended Data Fig. 3**

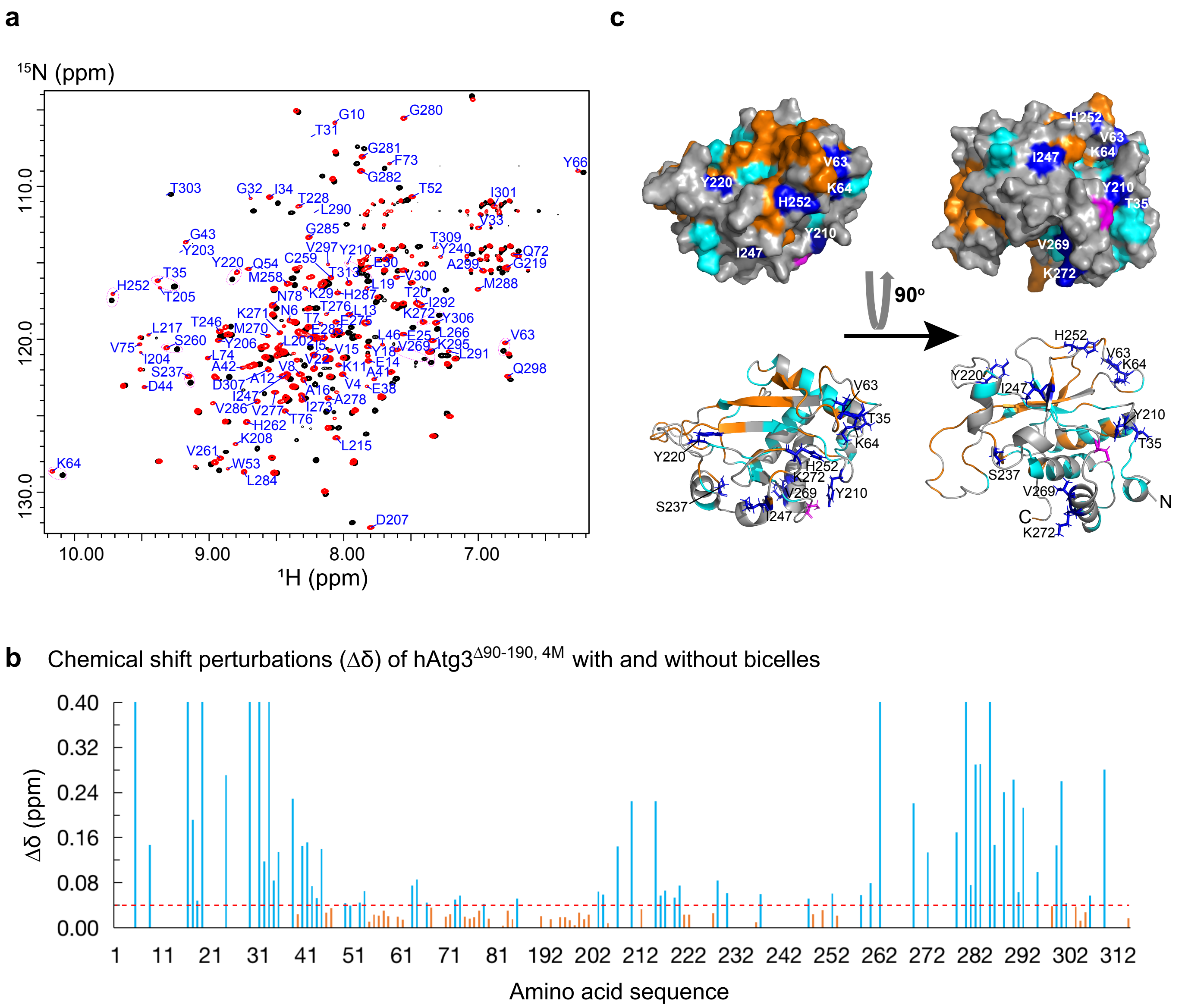


**Extended Data Fig. 4:** hAtg3$^{\Delta 90\text{-}190,\ 4M}$ binding to bicelles induces structural arrangements in its catalytic region.

**(a)** Overlay of $^{2}$H,$^{15}$N,$^{13}$C-labeled hAtg3$^{\Delta 90\text{-}190,\ 4M}$ TROSY spectra at pH 7.5 in the absence (black) and presence (red) of bicelles (DMPC:DMPG:DHPC = 4:1:20, molar ratio). Perturbed resonances are labeled with their assignments in bicelle-bound state.

**(b)** Plots of hAtg3$^{\Delta 90\text{-}190,\ 4M}$ $^{15}$N and $^{1}$H chemical shift perturbations (CSPs, Δδ) upon interaction with bicelles against residue numbers.

**(c)** Surface and ribbon representations of hAtg3 structure with perturbed residues (Δδ ≥0.04 ppm) colored light blue and unperturbed residues (Δδ < 0.04 ppm) colored orange. The active site C264 is highlighted in magenta and uncharacterized residues are colored grey. Some resonances that display large CSPs are circled in **(a)**; their corresponding residues are colored blue in **(c)**.

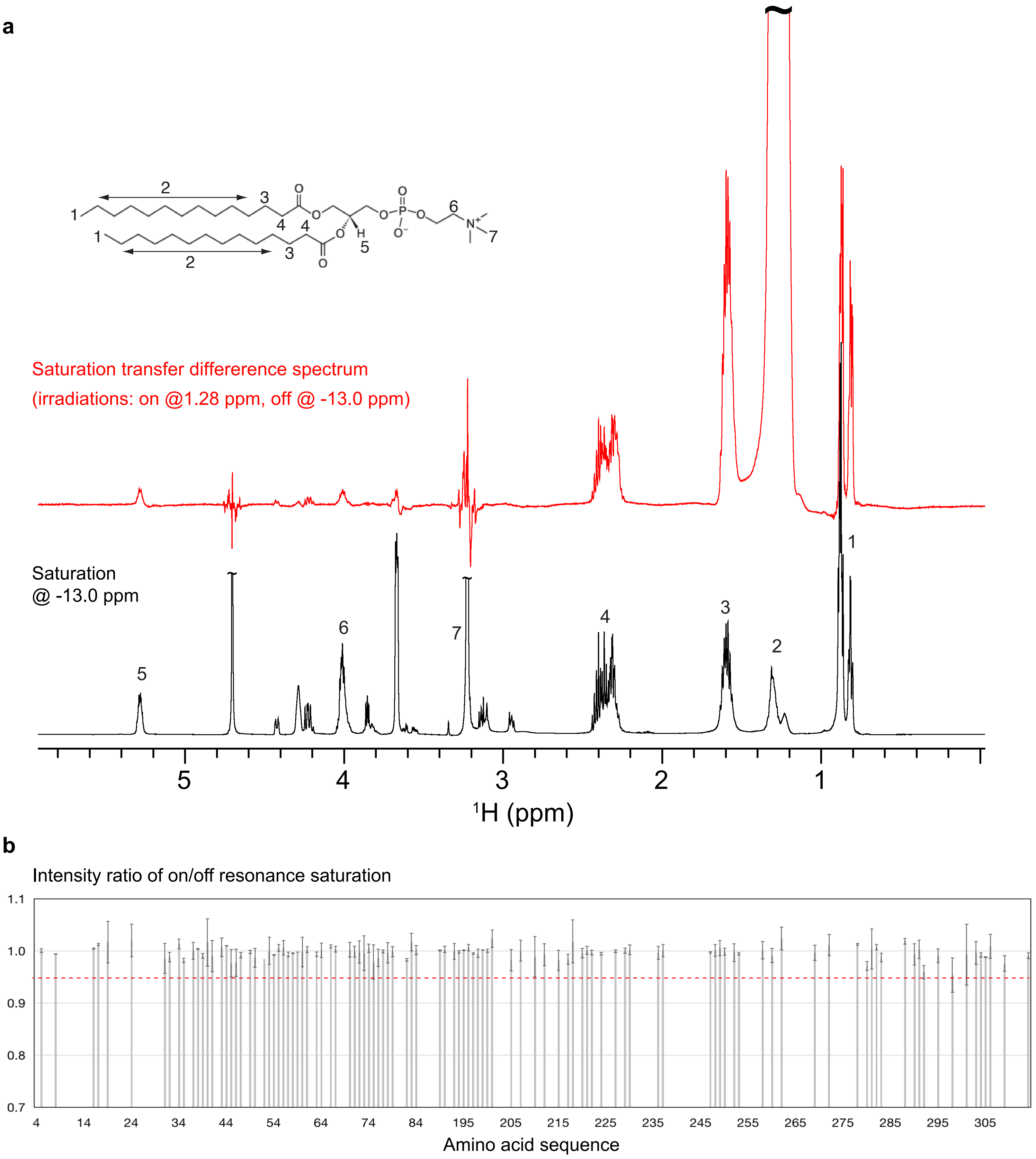


**Extended Data Fig. 5:** NMR cross-saturation experiments.

**(a)** Effective saturations of lipid resonances by irradiation at 1.28 ppm (peak #2).
Black: 1D $^{1}$H spectrum of $^{2}$H, $^{15}$N, $^{13}$C-hAtg3 in bicelles with saturation at -13.0 ppm (off).
Red: Saturation transfer difference spectrum of $^{2}$H, $^{15}$N, $^{13}$C-hAtg3 in bicelles.

**(b)** hAtg3$^{\Delta 90\text{-}190,\ 4M}$ in aqueous solution experiences few cross-saturation effects.
Plot of cross saturation effects against residue number for perdeuterated $^{15}$N, $^{2}$H- hAtg3$^{\Delta 90\text{-}190,\ 4M}$ in an 80% $D_2O$ and 20%$H_2O$ solution with saturations at -13.0 ppm (off) and 1.28 ppm (on).

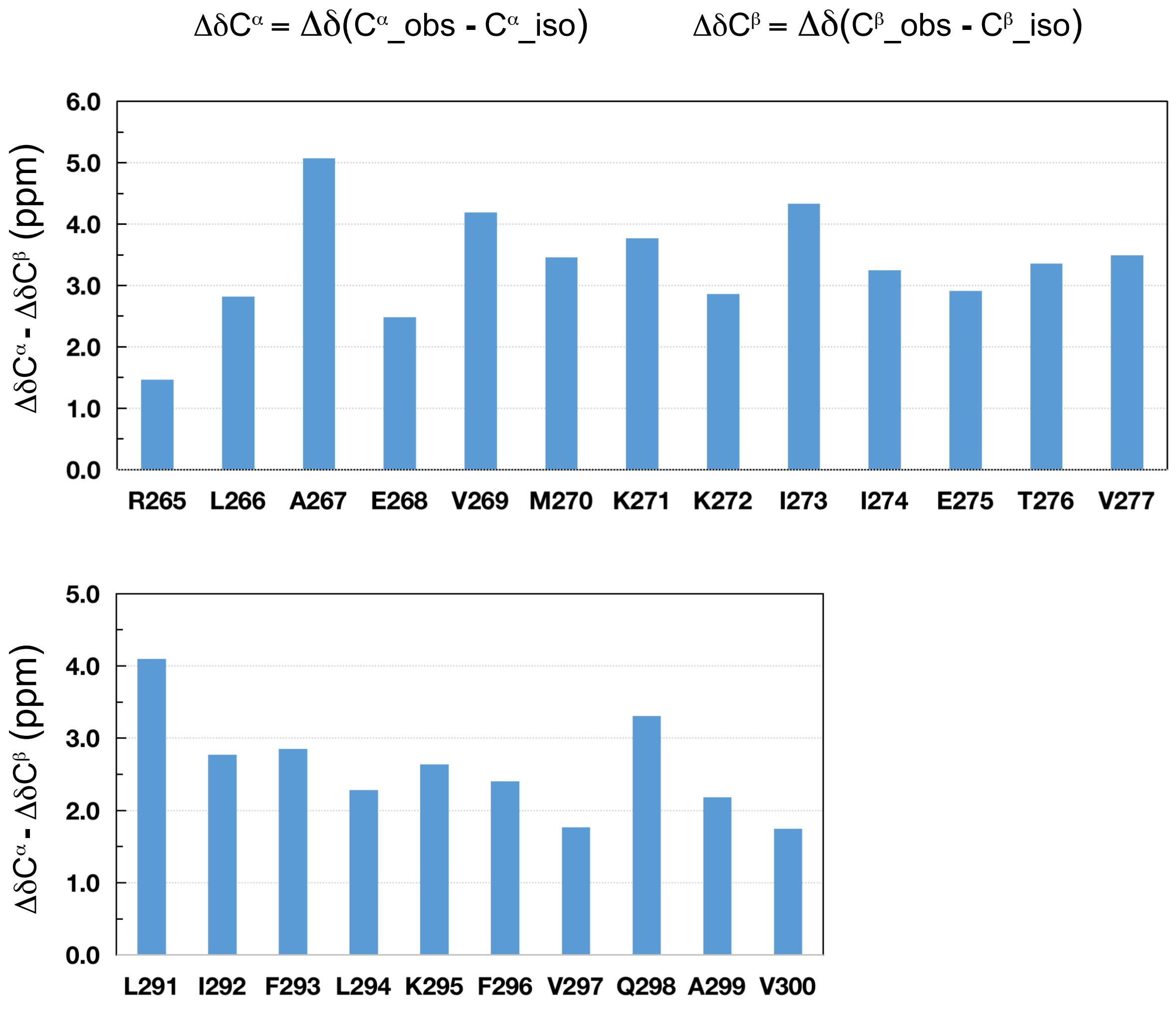


**Extended Data Fig. 6:** $^{13}C^{\alpha}$ and $^{13}C^{\beta}$ secondary chemical shifts indicate helical structures for residues 265 to 277 and residues 291 to 300 in bicelle-bound hAtg3$^{\Delta 90\text{-}190,\ 4M}$. $^{13}C^{\alpha}$_obs and $^{13}C^{\beta}$_obs are observed chemcial shifts while $^{13}C^{\alpha}$_iso and $^{13}C^{\beta}$_iso are isotropic values. Deuterium isotope shifts were not corrected.

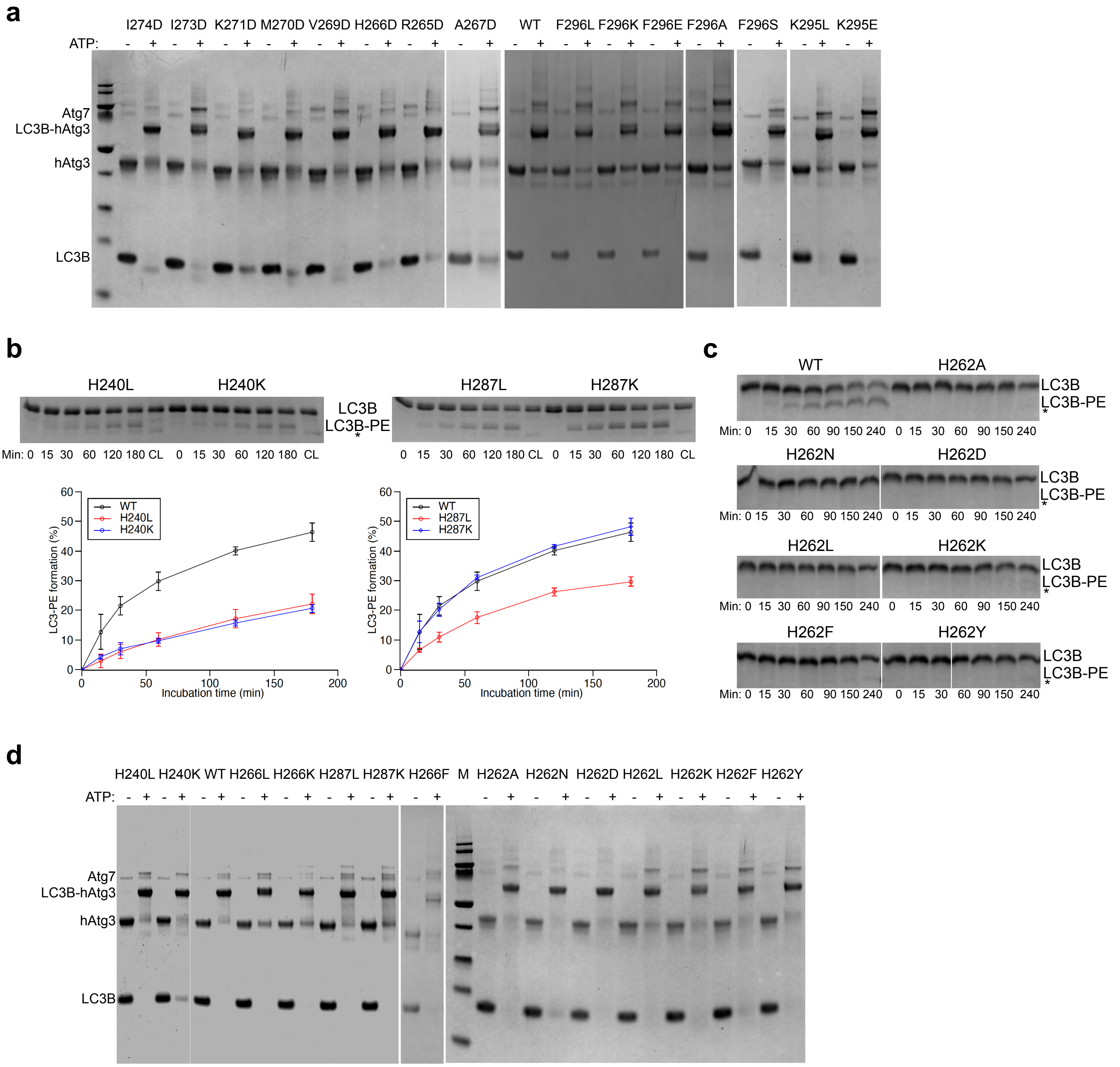


**Extended Data Fig. 7:** Gel analysis of hAtg3 mutations and their effects on LC3B-hAtg3 formation.

**(a)** NuPAGE (10% Bis-Tris, Invitrogen) images of hAtg3 WT and its mutants (5 μM) incubated with mouse Atg7 (0.5 μM), LC3B (5 μM) with (+) and without (-) ATP (1 mM) for 30 min at 37 °C. The formation of intermediate LC3B-hAtg3 is indicated. All experiments were repeated three times (n=3).

**(b)** Top: SDS-PAGE gel images of time dependent formation of LC3B-PE for H240L, H240K, H287L and H287K mutants. Bottom: Plots of time dependent formation of LC3B–PE for hAtg3 and mutants. Data are presented as mean ± SD. Quantification of conjugation reactions were obtained from three separate measurements.

**(c)** SDS-PAGE gel images of time dependent formation of LC3B-PE for hAtg3 and H262A, N, D, L, K, F, Y mutants. Asterisk indicates the degradation of LC3B in the presence of ATP.

**(d)** NuPAGE (10% Bis-Tris, Invitrogen) images of hAtg3 WT and its mutants (5 μM) incubated with mouse Atg7 (0.5 μM), LC3B (5 μM) with (+) and without (-) ATP (1 mM) for 30 min at 37 °C, pH 7.5. The formation of intermediate LC3B-hAtg3 is indicated. All experiments were repeated three times.

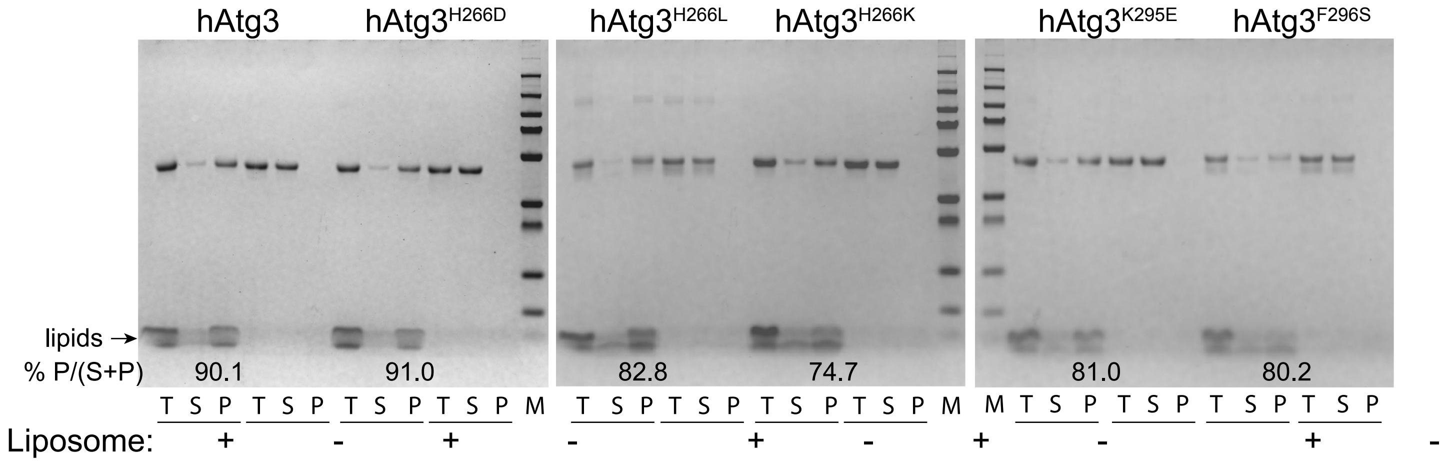


**Extended Data Fig. 8:** Interaction of hAtg3 and its mutants with liposomes in co-sedimentation assay. hAtg3 and its mutants (2 μM) were incubated with (+) and without (-) liposomes (800 μM) at 37 °C for 1 hr. Liposome-associated hAtg3 and its mutants were pelleted down by ultracentrifugation and analyzed by SDS-PAGE (10% Bis-Tris, Genscript). T: total; S: supernatant; P: pellet. The arrow indicates the lipid band.
